# A Harmonized Atlas of Spinal Cord Cell Types and Their Computational Classification

**DOI:** 10.1101/2020.09.03.241760

**Authors:** Daniel E. Russ, Ryan B. Patterson Cross, Li Li, Stephanie C. Koch, Kaya J.E. Matson, Ariel J. Levine

**Affiliations:** Division of Cancer Epidemiology and Genetics, Data Science Research Group, National Cancer Institute, NIH, Rockville, MD, USA; Spinal Circuits and Plasticity Unit, National Institute of Neurological Disorders and Stroke, NIH, Bethesda, MD, USA; Department of Neuroscience, Physiology and Pharmacology, Division of Biosciences, University College of London, London, UK

## Abstract

Single cell sequencing is transforming many fields of science but the vast amount of data it creates has the potential to both illuminate and obscure underlying biology. To harness the exciting potential of single cell data for the study of the mouse spinal cord, we have created a harmonized atlas of spinal cord transcriptomic cell types that unifies six independent and disparate studies into one common analysis. With the power of this large and diverse dataset, we reveal spinal cord cell type organization, validate a combinatorial set of markers for in-tissue spatial gene expression analysis, and optimize the computational classification of spinal cord cell types based on transcriptomic data. This work provides a comprehensive resource with unprecedented resolution of spinal cord cell types and charts a path forward for how to utilize transcriptomic data to expand our knowledge of spinal cord biology.

## INTRODUCTION

A revolution in single cell sequencing technologies is transforming many fields of biology. By sequencing the cDNA or open chromatin from many individual cells and using computational analysis to identify shared patterns of gene expression or epigenetic structure, we may simultaneously define cell “types”, characterize their molecular signatures, and track how each cell type in a tissue changes in different biological conditions such as development and disease. Within the central nervous system, this approach may also reveal the molecular basis of the impressive levels of neuronal diversity, can provide new marker genes for developing genetic tools to manipulate neuronal function, and may help to reveal the cellular basis of behavior.

In the postnatal mouse spinal cord alone, there have been nine papers profiling single cell RNA expression that, combined, cover a range of biological parameters, including age, tissue region, developmental lineage, and circuit features^1-9^. These studies provide a powerful and multi-faceted perspective on spinal cord cell types, yet despite this significant effort and a rich literature of spinal cord cell type characterization, there is still no consensus cell type “atlas” of the spinal cord. On the contrary, by conducting these studies independently, the number of nomenclature systems for spinal cord cell types has been multiplied without clarification of how these studies overlap, thereby leaving the underlying biology yet to be understood. Major obstacles include the lack of an accepted ground truth of cell types in this tissue^10^ that could form the basis of a reference atlas and the difficulty in comparing data between studies even when the same tissue types and techniques are used^3,5^. Indeed, these are among the “grand challenges” that scientists face as we re-discover the cells and tissues we study through the perspective of single cell profiling^11^.

To begin to overcome these challenges within the mammalian central nervous system, we sought (1) to establish a harmonized atlas of postnatal spinal cord cell types that is shared across biological time, experimental technique, and laboratory, (2) to enhance the usability of this data for broader field of spinal cord biology, and (3) to test different tools to facilitate the future classification of cells into these types. We began by performing an integrated and merged analysis of the raw data from the first six publicly available postnatal spinal cord single cell datasets. Next, we clustered the cells and nuclei of this meta-dataset to reveal 15 non-neural and 69 neural cell types, thereby providing a cell type resolution and characterization that surpasses all prior studies. By analyzing gene expression profiles across families of clustered cell types, we created a combinatorial panel of marker genes and validated it with high-content in situ hybridization. Finally, we tested a range of automated classification algorithms and identified a two-tiered model based on label transfer and neural networks as the best method for classifying spinal cord cell types. We have now developed “SeqSeek”, a web-based resource for querying this data by gene or cell type and for accessing automated classification algorithm of any spinal cord cell or nucleus from raw sequencing data.

## RESULTS

### Merged Analysis of Spinal Cord Cells and Nuclei

The work here is based on a merged dataset with over one hundred thousand cells and nuclei from the first six published studies of the postnatal mouse spinal cord^1-6^. These studies cover a range of biological and experimental parameters (Figure 1A and Supplemental Figure 1). To best compare the data from these studies, we began with the raw sequencing reads from each study and performed our own data processing with uniform methods and filters. All sequencing reads were aligned to a common genomic sequence that included both exons and introns and common filtering thresholds were used for inclusion (>200 genes per cell/nucleus) and exclusion (<5% percent of genes from mitochondria). As a result, this merged dataset contains more cells and nuclei than were analyzed in the original studies and a uniform set of genes (Supplemental Figure 1).

**Figure 1.**
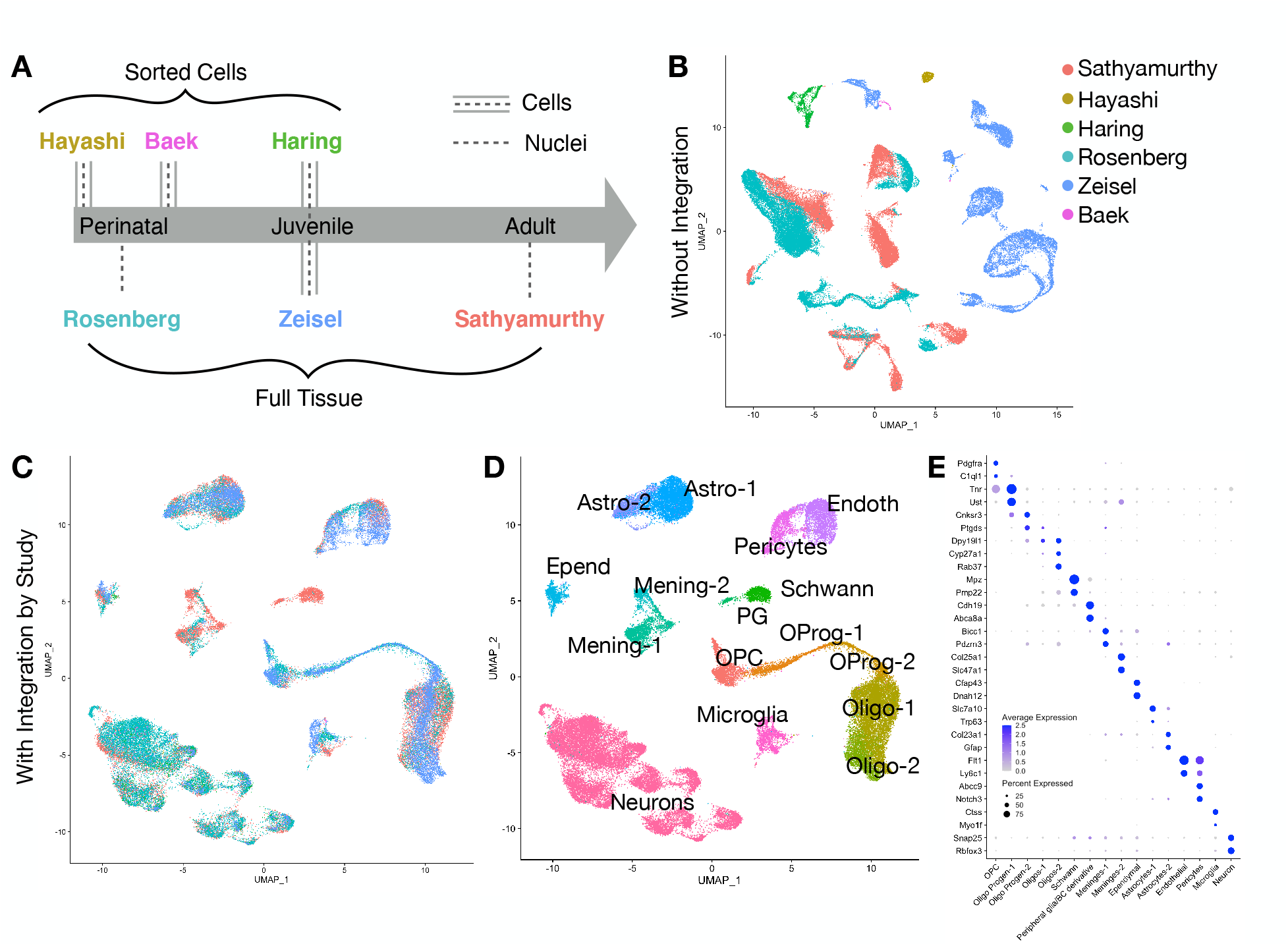
Integration of six independent studies on single cell spinal cord data reveals the major cell types of the spinal cord. (A) Six independent studies that used single cell/nucleus RNA sequencing to analyze mouse spinal cord cell types were analyzed, covering a range of mouse ages and technical approaches. (B) UMAP presentation of the 52,623 cells/nuclei in the final dataset, without integration and colored by the study of origin (colors in the legend). (C) UMAP presentation of the same 52,623 cells/nuclei in the final dataset, integrated by study and colored by the study of origin (same colors as in (B)). (D) UMAP presentation of the cells/nuclei in the final dataset, integrated by study and colored by cell type. (E) Dot plot of the expression of marker genes for the major coarse cell types. Average expression for each cluster is shown by color intensity and the percent of cells/nuclei in each cluster that expressed each gene is shown by dot diameter.

Our first major goal was to create a harmonized atlas of the major spinal cord cell types that are shared across these studies. Previous reports have used the correlation in gene expression between clusters to link cell types across studies, but this approach yielded weak correlations, even between studies in which the same sample age and tissue dissociation method were used^3,5^. We hypothesized that co-clustering cells and nuclei across all of the studies would provide an improved ability to relate cell types in one study to those in another. We performed dimensionality reduction using principal component analysis and visualized the cells and nuclei using UMAPs. Unfortunately, the cells or nuclei from each study segregated from each other almost completely, indicating that the study of origin is a major source of variability in the dataset (Figure 1B). This technical limitation obscured all cell type distinctions.

To reduce experimental sources of variability and reveal the core set of spinal cord cell types, we used a recently developed integration method to align the cells and nuclei across studies ^12-15^. With this approach, the cells and nuclei from all six studies were spatially interposed in a UMAP visualization of principal component space (Figure 1C) and separated into groupings that each expressed a panel of well-established cell type markers such as Snap25 (neurons), Mbp (oligodendrocytes), Aqp4 (astrocytes), and Ctss (microglia). After preliminary clustering and the removal of low-quality clusters and doublets (see Methods), we obtained a merged dataset of over fifty thousand cells and nuclei. The majority of these cells/nuclei from this analysis are from the three studies that used high throughput collection and barcoding techniques (the Sathyamurthy, Rosenberg, and Zeisel datasets) (Supplemental Figure 1). A comparison across studies revealed that these high throughput studies detected fewer genes per cell/nucleus than studies that used single well technical approaches (the Hayashi, Haring, and Baek datasets), and studies that used cells (the Hayashi, Haring, Zeisel, and Baek datasets) detected more genes per cell/nucleus but had relatively higher levels of immediate early gene and stress gene expression than did studies that used nuclei (the Sathyamurthy and Rosenberg datasets) (Supplemental Figure 1). These trends across technical approaches were expected based on other reports (reviewed^12^).

### A Harmonized Atlas of Major Cell Types

Next, we performed coarse clustering to define the major cell types of the mouse spinal cord (Figure 1D,E). Sixteen major types were identified that represent all known classes of spinal cord cell types; a characterization and resolution that surpasses all of the original six studies in capturing the full diversity of spinal cord cell types. These cell types are: (1) oligodendrocyte precursor cells; (2-3) two stages of oligodendrocyte progenitors; (4-5) two types of oligodendrocytes that likely correspond to myelinating and mature cell types and that blend into each other; (6) Schwann cells; (7) peripheral glia; (8-9) two types of meninges that likely correspond to vascular leptomeningeal cells and arachnoid barrier cells; (10) ependymal cells that surround the central canal; (11-12) two types of astrocytes that likely correspond to a major population of regular astrocytes and a minor population of Gfap-expressing proliferating/activated/white matter astrocytes; (13-14) two types of vascular cells that likely correspond to endothelial cells and pericytes; (15) microglia; and (16) neurons, which are discussed in detail below.

As expected, the cell types that were derived from each study corresponded to the techniques used to isolate the cells or nuclei (Supplemental Figure 1). The three studies that FACS sorted neurons from the spinal cord (Hayashi, Haring, and Baek datasets) predominantly gave rise to cells in the neuronal sub-clusters as well as the non-neural cells most likely represent doublets. Moreover, among the three studies that examined all cell types, the early postnatal Rosenberg study showed an enrichment of immature cells of oligodendrocyte lineage relative to the adult Sathyamurthy study, while the adolescent Zeisel study showed an intermediate distribution. The only study to dissect the spinal cord including the dorsal and ventral spinal roots (the Sathyamurthy dataset) was the only source of Schwann and peripheral glia cells that would be located in these roots.

### A Harmonized Atlas of Neuronal Populations

We next focused our analysis on neuronal populations to further probe their impressive diversity and to define a reference set of cell types for understanding the spinal cord cellular basis of behavior. Based on the coarse cell type assignments above, we selected and clustered all neuronal cells/nuclei. Preliminary analysis revealed that putative dorsal horn clusters separated well in principal component space while putative mid and ventral horn clusters did not, which prompted us to perform a targeted sub-clustering of all mid and ventral cells/nuclei (see Methods). 69 neuronal clusters were identified (Figure 2A, Table 1, Supplemental Movie 1, Supplemental Table 2) and the neurotransmitter status and putative regional location (dorsal horn, mid region, ventral horn) were determined by marker gene expression and comparison to the original six studies. We observed 20 dorsal excitatory clusters, 14 dorsal inhibitory clusters, 10 deep dorsal/mid excitatory clusters, 7 deep dorsal/mid inhibitory clusters, 8 ventral excitatory clusters, 6 ventral inhibitory clusters, 3 cholinergic motoneuron clusters, and 1 cluster of cerebrospinal fluid contacting neurons. As was observed in the full dataset with all cell types, neuronal cells/nuclei from studies that used massively parallel approaches (Sathyamurthy, Rosenberg, Zeisel) had fewer genes per cell/nucleus and that studies that those which used nuclei (Sathyamurthy and Rosenberg) had lower levels of immediate early gene and stress gene expression than studies that used cells (Hayashi, Haring, Zeisel and Baek) (Supplemental Figure 2).

**Table 1.**
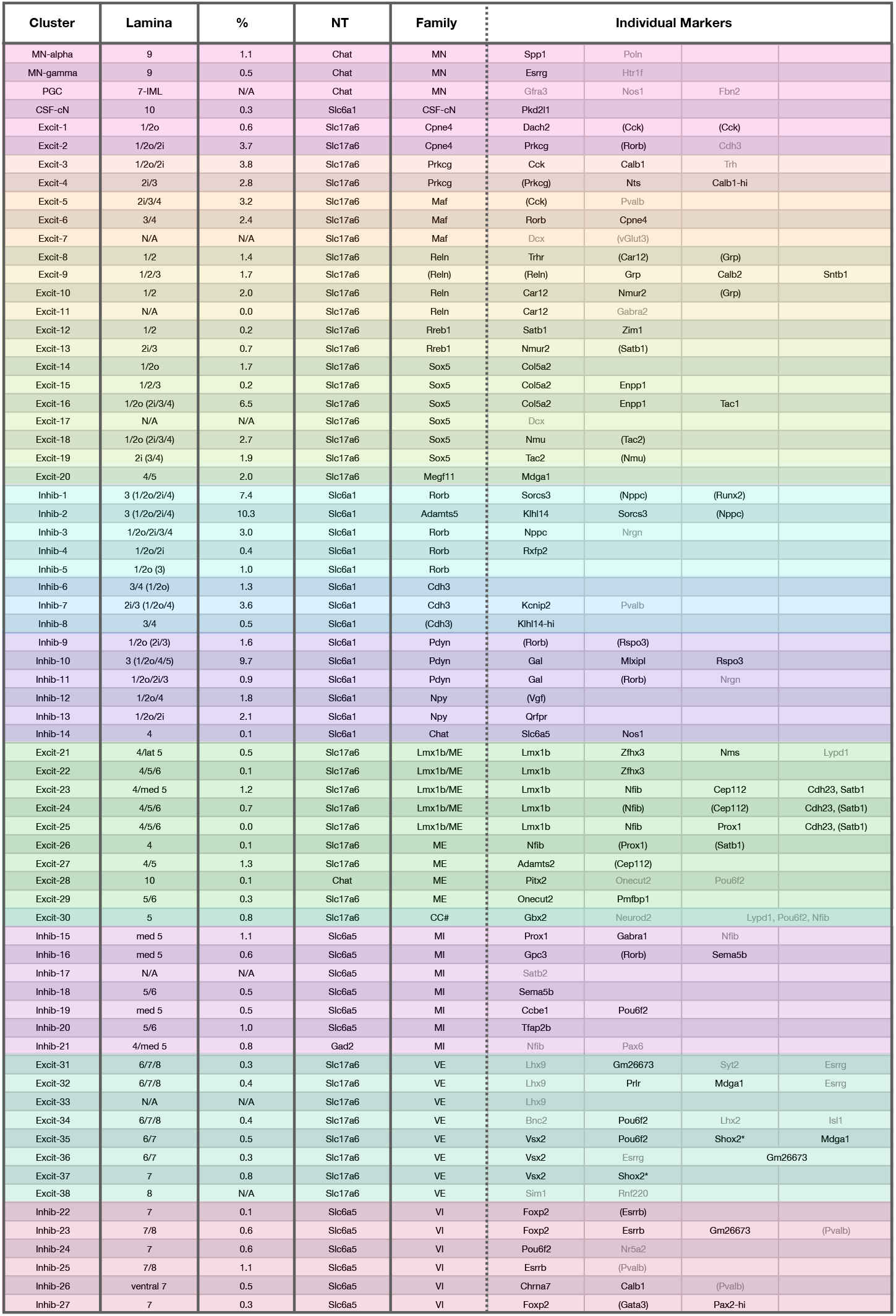
Cell-type census of 69 populations of spinal cord neurons. The lamina, prevalence, a neurotransmitter marker gene, “family” and individual markers for each neuronal cluster are shown. The clusters are color coded to correspond approximately to their color in Figure 2A. The prevalence of each cluster was determined by counting the confidently assigned cells of each type based on RNA in situ hybridization on sections from three animals and are presented as the percent of the total number of confidently assigned neurons. Genes in parenthesis are expressed at lower levels. Genes in gray were not validated (due to probe failure, being present only in postnatal animals, or were not included in the analysis). ^#^ denotes a putative identity (see main text). * denotes a marker that was validated using RNAScope V2 but did not work in the RNAScope Hiplex assay.

**Figure 2.**
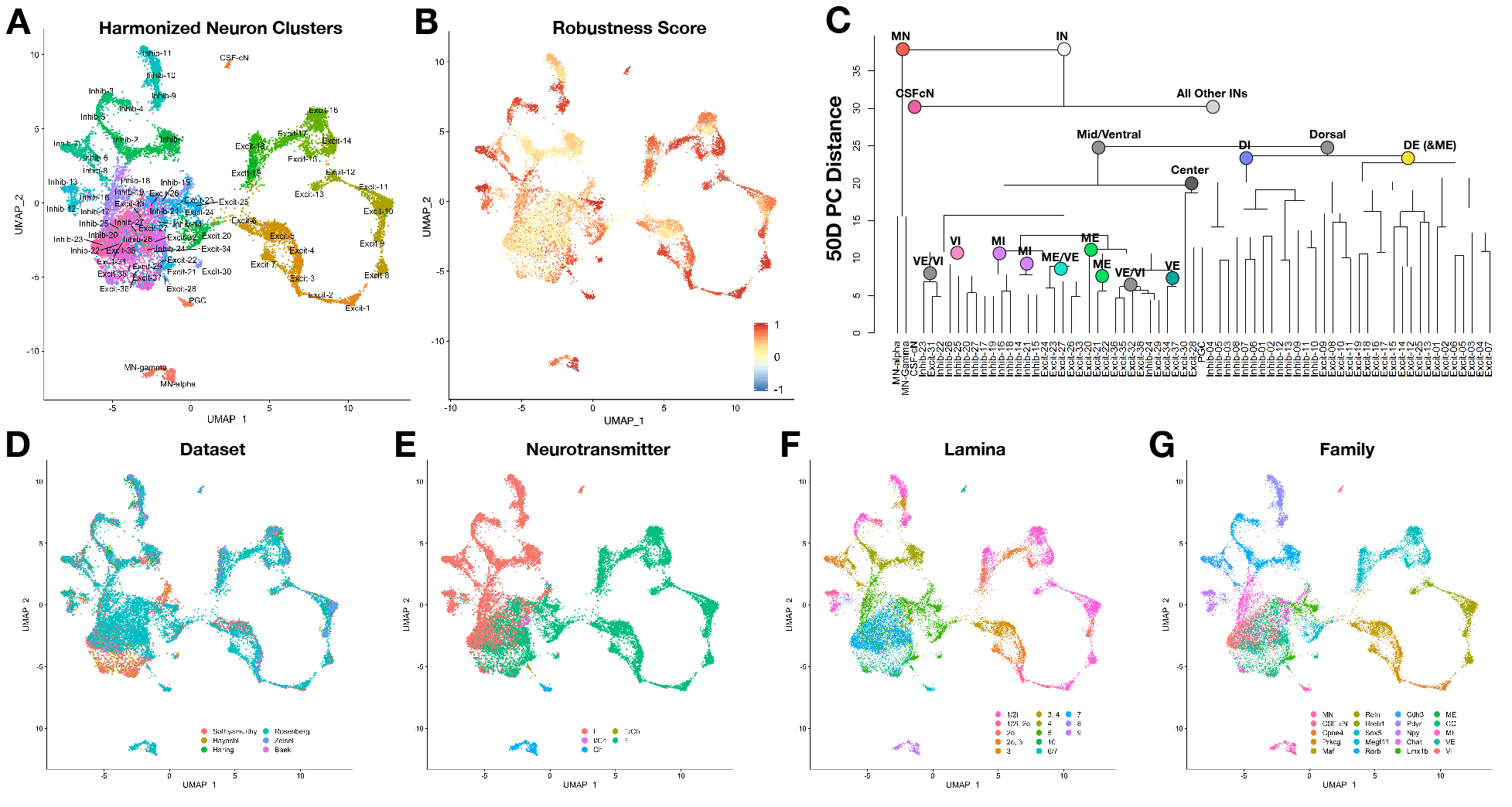
Harmonized atlas of 69 populations of spinal cord neurons. (A) UMAP presentation of 19,353 neuronal cells/nuclei of the postnatal mouse spinal cord, colored and annotated by cell-type cluster. (B) The same cells/nuclei, colored by robustness (silhouette) score, which was calculated based on bootstrapped co-clustering frequency (see Methods). (C) Dendogram showing the relationships between the 69 neuronal cell types based on their distance from each other in the 50-dimensional principal component (PC) space. MN=motoneuron; IN=interneurons (and projection neurons); CSF-cN=cerebrospinal fluid contacting neurons; DE=dorsal excitatory; DI=dorsal inhibitory; ME=mid excitatory; MI=mid inhibitory; VE=ventral excitatory; VI=ventral inhibitory; “center” represents a group of 3 cell types located near lamina X – the center of the spinal cord. (D-G) UMAP presentation of 19,353 neuronal cells/nuclei of the postnatal mouse spinal cord, colored by study of origin (E), neurotransmitter (F), lamina (G), and family (H). (E) I=inhibitory, I/Ch=inhibitory cholinergic, Ch = cholinergic; E/Ch=excitatory cholinergic; E=excitatory. (F) Laminae were assigned based on in situ hybridization validation experiments and are colored by the approximate depth from the dorsal surface of the cord (hot pink to violet). (G) See main text for description of neuronal families.

To determine the robustness of these clusters, we used a bootstrapped co-clustering test of the consistency with which cells and nuclei in each cluster remain together upon repeated clustering (Figure 2B, Supplemental Figure 2). As expected, dorsal clusters showed very high robustness with this measure, whereas mid and ventral clusters showed moderate to low robustness, a general feature that was consistent with previous observations^1,4^. This most likely reflects the highly similar and even overlapping patterns of gene expression amongst mid and ventral clusters. Similarly, a dendrogram analysis of the distance between the clusters within the 50-dimensional principal component space also revealed that dorsal clusters were well separated from each other, while mid and ventral clusters were much closer to each other in this reduced gene expression space (Figure 2C). Intriguingly, neurons that are located at the spatial mid-point between the dorsal and ventral sides of the cord (preganglionic cells and two excitatory populations near the central canal) were organized as a single branch (Figure 2C; “center”), further underscoring the importance of spatial distribution as an organizing principle in the spinal cord.

Next, we sought to characterize these clusters at a molecular level and to define their marker genes. There are multiple approaches for identifying cell type markers based in single cell data. Commonly used methods such as such as the Wilcox Rank Sum test and ROC analysis use differential expression to identify genes that are enriched within one identified cell cluster as compared to all other clusters and we used this approach to generate candidate markers for each cluster (Supplemental Table 1). However, these approaches do not prioritize markers that are shared between related clusters or those markers that are well-established for a given tissue, nor do they produce an efficient final set of markers that can be used to define all neuronal cell types. To overcome these obstacles, we therefore used a combination of Wilcox and ROC individual cluster markers, Wilcox and ROC markers for dendrogram branches, and established markers from the literature to generate a panel of combinatorial markers for spinal cord neurons that follows a “family name” and “given name” analogy. For example, Excit-14 through Excit-19 comprise the “Sox5” family. They are distinguished by expression of Col5a2 (Excit-14), Col5a2 and Enpp1 (Excit-15), Col5a2, Enpp1, and Tac1 (Excit-16), Dcx expression and being present almost exclusively at early post-natal stages (Excit-17), Nmu (Excit-18), and Tac2 (Excit-19) (Figure 3 and Table 1).

**Figure 3.**
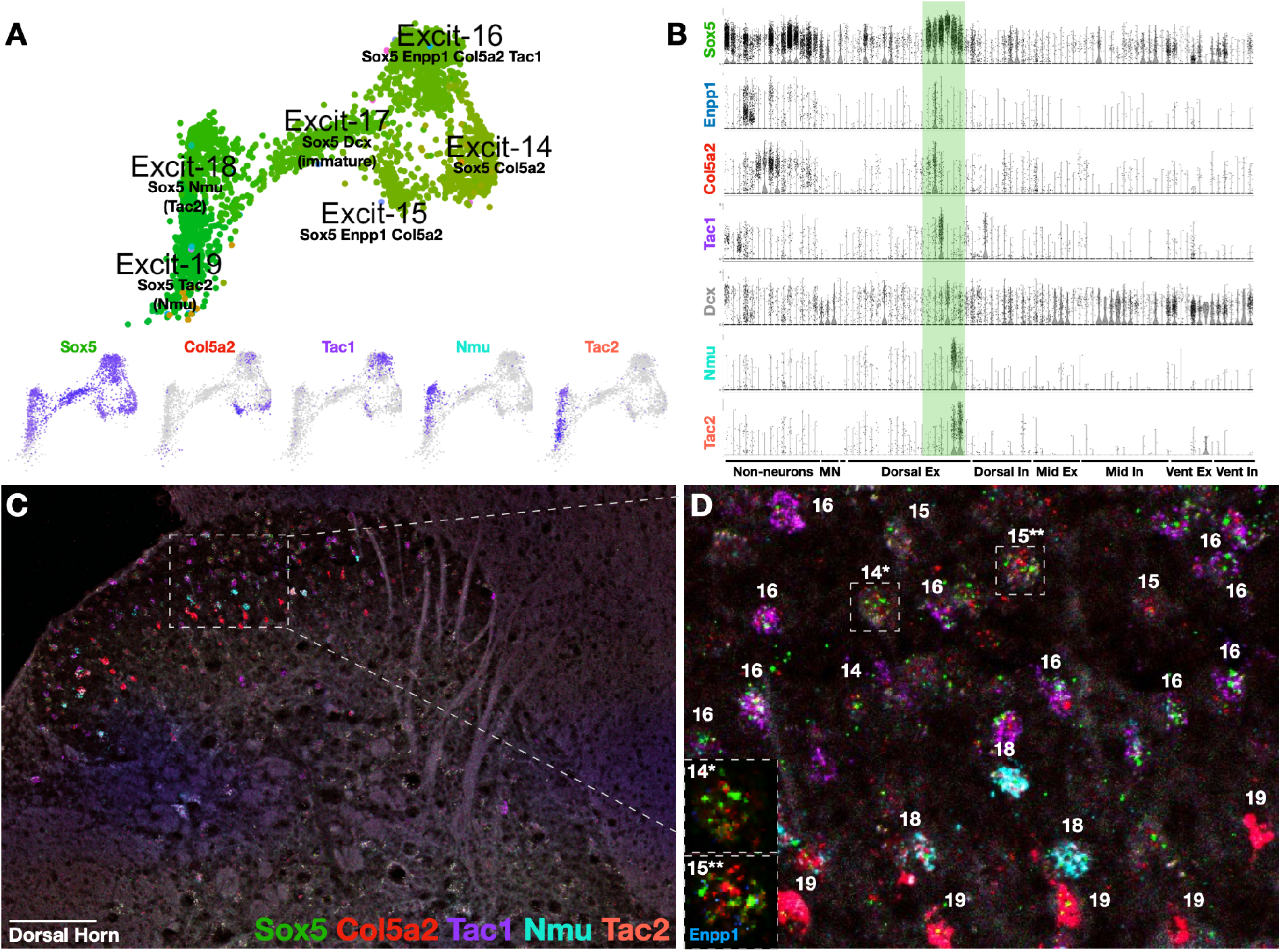
The Sox5 dorsal excitatory family is sub-divided into individual clusters by a panel of marker genes. (A) The region of the neuron cell types UMAP for Excit-14 through Excit-19, labeled with relevant marker genes (top) and single feature plots of selected marker genes, where expression is coded from absent (light gray) through highly expressing (dark purple) (bottom). (B) Violin plot of the distribution of selected marker genes across all 84 clusters, including non-neurons, MN=motoneurons, CSF-cN=cerebrospinal fluid contacting neurons, DE=dorsal excitatory; DI=dorsal inhibitory; ME=mid excitatory; MI=mid inhibitory; VE=ventral excitatory; VI=ventral inhibitory. Each dot represents a single cell or nucleus and the Sox5 family dorsal excitatory family is highlighted with the olive green bar. (C) RNA in situ hybridization of selected marker genes Sox5, Col5a2, Tac1, Nmu, Tac2 on an adult mouse lumbar spinal cord section. 20x tiled image, with brightness and contrast adjusted. (D) Zoomed region of (C). Cells were assigned to individual excitatory clusters (see individual numbers) based on marker gene expression. Inset show representative cells of Excit-14 (14*) and Excit-15 (15**) with in situ hybridization for Sox5 (green), Col5a2 (red), Enpp1 (blue). Scale bar is 100 μm.

To determine whether this panel of markers corresponded to in situ gene expression patterns and to define the anatomical distribution of each cluster, we performed high-content in situ hybridization with combinatorial sets of marker gene probes (Supplemental Table 3). We tested 95 unique genes (of which 79 showed reliable expression in the adult lumbar spinal cord) and analyzed gene expression in ten overlapping sets of 12 genes each. For each set, hundreds of cells were counted from three spinal cords and their locations mapped by lamina. Using this approach, 71% of neurons in the adult lumbar spinal cord could be identified as belonging to one of the 69 neuronal clusters (2057/2894 total) and an additional 9% of neurons could be identified as belonging to pairs of closely related clusters (266/2894 total) (Supplemental Table 3). We found that some sets (such as those that sub-type dorsal inhibitory neurons) could be used to identify 80-90% of cells, while other sets (such as those that sub-type mid and ventral neurons) could only identify 40-60% of neurons. This further supports the view that dorsal neurons are more molecularly distinct while mid and ventral neurons are more difficult to distinguish from one another. This detailed in situ hybridization analysis also revealed the in-tissue location and prevalence of each of the lumbar adult neuronal cell types and can serve to translate single cell sequencing data back into tissue-based analysis.

The cell type markers, laminar distribution, and estimated prevalence of each cluster are shown in Table 1, Figure 3, and Supplemental Figure 3 and are presented by family, with comments, as follows.

#### MN (3 clusters)

The motoneuron (MN) family includes alpha motoneurons (MNa) which had relatively higher levels of Spp1 and Poln, gamma motoneurons (MNg) which had relatively higher levels of Esrrg and Htr1f, and the related preganglionic cells (PGC) which expressed Gfra3 and Nos1. This family was only comprised of nuclei from the Sathyamurthy and Rosenberg datasets, although the Zeisel dataset was also expected to include motoneurons. Of note, we did not detect refined sub-populations of MNa or PGC, although it is likely that further work will sub-fractionate MNa into fast and slow populations, or even specific muscle pools. Motoneurons are the final output cell through which the central nervous system controls muscles and the autonomic system and can be found in lamina 9 (MNa and MNg) or lamina 7/intermediolateral nucleus (PGC). Supplemental Figure 3A.

#### CSF-cN (1 cluster)

<u>C</u>erebro<u>s</u>pinal fluid <u>c</u>ontacting <u>n</u>eurons were distinguished by Pkd2l1, as well as Pkd1l2. This cluster was very distinct from other neuronal populations, inhibitory, and also expressed the early neuron marker Sox2 and the V2b lineage markers Gata2 and Gata3, suggesting an “immature” phenotype. Supplemental Figure 3A.

#### Dorsal Excitatory

##### Cpne4 (2 clusters)

This dorsal, excitatory family was comprised of Excit-1 and Excit-2. Excit-1 was a rare subset, both in the harmonized clusters and in the in situ counts, that also expressed Dach2 and Excit-2 was more prevalent and co-expressed Prkcg as well as Cbln2. Supplemental Figure 3B.

##### Prkcg (2 clusters)

This dorsal, excitatory family was comprised of Excit-3 and Excit-4. Prkcg is a classic marker gene in the spinal cord and defined this family together with the neuropeptides Cck and Trh (Excit-3) and Nts (Excit-4). Both subsets also expressed Calb1, although it was not specific to these clusters. This family was also close to Excit-7, an immature cluster grouped with the Maf family. Supplemental Figure 3B.

##### Maf (3 clusters)

This dorsal, excitatory family was comprised of Excit-5, Excit-6, and Excit-7. All three clusters expressed enriched levels of Rora (which was broadly expressed in many other clusters at lower levels). Excit-5 also expressed Pvalb, Excit-6 expressed Rorb and Cpne4, and Excit-7 was distinguished by having only nuclei from the Rosenberg dataset and expressed the immature neuron marker Dcx, suggesting an immature phenotype. The similarity of Excit-7 with Excit-3, Excit-4, Excit-5, and Excit-6 suggests a shared lineage relationship between these families. This family also expressed low levels of Slc17a8 (vGlut3). Supplemental Figure 3B.

##### Reln (4 clusters)

This dorsal, excitatory family was comprised of Excit-8, Excit-9, Excit-10, and Excit-11. These clusters expressed enriched levels of Car12 (in particular in Excit-9 and Excit-10), the neuropeptide receptors Trhr (Excit-8), Npr1 (Excit-9 and Excit-10), and Nmur2 (Excit-10) and the neuropeptide Grp (Excit-9). Supplemental Figure 3C.

##### Rreb1 (2 clusters)

This dorsal, excitatory family was comprised of Excit-12 and Excit-13. These clusters also express Satb1 and either Zim1 (Excit-12) or Nmur2 and Crh (Excit-13). Supplemental Figure 3C.

#### Sox5 (6 clusters)

This dorsal, excitatory family was comprised of Excit-14, Excit-15, Excit-16, Excit-17, Excit-18, and Excit-19. Within this family, Excit-14 and Excit-15 were slightly separated and also similar to the Rreb1 family clusters and expressed Col5a2 (Excit-14) or Col5a2 and Enpp1 (Excit-15). Excit-16, Excit-18, and Excit-19 expressed the neuropeptides Tac1 (Excit-16), Nmu-hi/Tac2-lo (Excit-18), and Tac2hi/Nmu-lo (Excit-19). Excit-17 included almost exclusively nuclei from the Rosenberg dataset and expressed the immature neuron marker Dcx, suggesting an immature phenotype. As this cluster was similar to Excit-16, Excit-18, and Excit-19, this may suggest a shared lineage relationship between these clusters. Figure 3.

##### Megf11 (1 cluster)

This Excit-20 cluster displayed features of dorsal excitatory neurons and mid excitatory neurons, being located in lamina 4/5 and being grouped with mid neurons in principal component space in the uMAP and dendogram analysis. It expressed Megf11 and Mdga1.

#### Dorsal Inhibitory

##### Rorb & Adamts5 (5 clusters)

This dorsal, inhibitory family was comprised of Inhib-1, Inhib-2, Inhib-3, Inhib-4, and Inhib-5. Each of these clusters, except Inhib-2, expressed Rorb. Inhib-2 is grouped with this family based on its proximity in principal component space, as reflected in the uMAP and dendogram analysis. In addition to Rorb, Inhib-1 expressed Sorcs3, Inhib-3 expressed Rorb and Nppc as well as as Nrgn, Inhib-4 expressed Rorb and Rxfp2, and Inhib-5 did not express these other genes. Inhib-2 expressed Sorcs3 and Adamts5. Inhib-1 and Inhib-2 represent deeper dorsal (lamina 3) clusters, Inhib-3 was distributed throughout the dorsal horn, and Inhib-4 and Inhib-5 were relatively rare clusters (as judged by the harmonized cluster sizes and the in situ counts) and were found in the superficial laminae (1/2). Supplemental Figure 3D.

##### Cdh3 (3 clusters)

This dorsal, inhibitory family was comprised of Inhib-6, Inhib-7, and Inhib-8. Inhib-6 and Inhib-7 expressed Cdh3 and were distinguished by co-expression of Kcnip2 and Pvalb in Inhib-7. While Inhib-8 contained only low levels of Cdh3 in this analysis, Cdh3 expression was confirmed by in situ hybridization and this cluster was included in this family based on proximity in principal component space as reflected in the uMAP and dendogram analysis. Inhib-8 expressed Klhl14. Supplemental Figure 3D.

##### Pdyn (3 clusters)

This dorsal, inhibitory family was comprised of Inhib-9, Inhib-10, and Inhib-11. Each of these clusters expressed Pdyn, while Inhib-10 also expressed Gal and Mlxipl and Inhib-11 also expressed Gal only. Of note, the clusters in this family also expressed Rorb and Nrgn. Supplemental Figure 3E.

##### Npy (2 clusters)

This dorsal, inhibitory family was comprised of Inhib-12 and Inhib-13. These clusters expressed Npy and were distinguished by low levels of Vgf (Inhib-12) or by expression of Qrfpr (Inhib-13). Supplemental Figure 3E.

##### Chat (1 cluster)

This Inhib-14 cluster is a deep dorsal (lamina 4), inhibitory and cholinergic population and also expressed Nos1.

#### Mid/Deep Dorsal Horn Clusters

Of note, mid clusters generally were less robust than dorsal clusters.

##### Excitatory (ME)/Lmx1b (5 clusters)

This family of mid, excitatory clusters was comprised of Excit-21, Excit-22, Excit-23, Excit-24, and Excit-25. These clusters expressed Lmx1b, suggesting a dI5/dIL^B^ embryonic origin. All of the clusters except Excit-25 expressed Tacr1 and Excit-21 also expressed Lypd1, suggesting that these are candidate ascending populations^3^. These clusters could also be distinguished by expression of Zfhx3 (Excit-21 and Excit-22) or Nfib (Excit-23, Excit-24, and Excit-25), which corresponded to lateral Zfhx3 and medial Nfib sub-types. Other markers sub-divided the clusters in a combinatorial manner, including Nms (Excit-21), Bcl11a (Excit-22 through Excit-25), Satb1 and Cdh23 (Excit-23, Excit-24, and Excit-25), Cep112 (Excit-23 and Excit-24), and Prox1 (Excit-25). Of note, nearly all of the cells and nuclei in this family were from the Rosenberg and Sathyamurthy datasets. Supplemental Figure 3F.

##### Excitatory (ME) (4 clusters)

This family of mid, excitatory clusters was comprised of Excit-26, Excit-27, Excit-28, and Excit-29. These clusters do not express Lmx1b, in contrast to the other mid excitatory family and may be derived from ventral embryonic lineages. Excit-26 expressed Nfib, Excit-27 expressed Adamts2, Excit-28 expressed Chat and Pitx2 and thus likely corresponds to V0c neurons, and Excit-29 expressed Pmfbp1. Excit-28 and Excit-29 also express Onecut2 and Pou6f2, potentially revealing a link with ventral cell types. Of note, nearly all of the cells and nuclei in this family were from the Rosenberg and Sathyamurthy datasets and Excit-26 in particular was predominantly from the Rosenberg dataset. Supplemental Figure 3A and 3F.

##### Excit-30/CC^#^ (1 cluster)

This cluster was marked by Gbx2, Neurod2, and Sp8 and there was partial evidence that it corresponded to Clarke’s column. This cluster expressed multiple genes associated with Clarke’s column including Chmp2b, Syt4, Ebf3, Rgs4, and Enc1^6^. The Clarke’s column marker gene, Gdnf, was expressed at very low levels in the merged dataset, but was present in several Excit-30 cells. However, this cluster only contained two defined spinocerebellar cells from the Baek et al. dataset while the majority of this cluster was from the Hayashi dataset, arguing against a Clarke’s column identity and also suggesting a V2 embryonic lineage. As the in situ hybridization experiments were performed on lumbar spinal cord sections, we did not validate markers for this cluster.

##### Inhibitory (MI) (7 clusters)

This family of mid, inhibitory clusters was comprised of Inhib-15, Inhib-16, Inhib-17, Inhib-18, Inhib-19, Inhib-20, and Inhib-21, all of which expressed the glycinergic marker Slc6a5 (with the exception of Inhib-21) and also the gabaergic marker Gad2. Inhib-15 expressed Prox1, Gabra1, and Nfib, Inhib-16 expressed Gpc3 and Sema5b, Inhib-17 expressed Satb2, Inhib-18 expressed Sema5b, Inhib-19 expressed Ccbe1 and Pou6f2, Inhib-20 expressed higher levels of Tfap2b as well as Zfhx3, and Inhib-21 expressed Nfib and was distinguished by having only Gad2 and not Slc6a5 and was mainly derived from the Rosenberg dataset. Supplemental Figure 3G.

#### Ventral Clusters

In general, the ventral clusters had less distinct gene expression patterns and were less robust than dorsal and mid clusters; therefore, the final identities of these clusters should be considered with caution. We identified several genes that contribute to overlapping gene expression patterns across clusters by being present in a spatial region of the cord and in diverse mid/ventral cell types. For example, Pou6f2 was expressed in the deep dorsal horn and in the dorsal part of the ventral horn and was enriched in mid-excitatory (Excit-21, Excit-28, and Excit 30), ventral excitatory (Excit-34 and Excit-35), and a ventral inhibitory (Inhib-24) clusters that are located within this domain. Similarly, Nfib was expressed in the medial deep dorsal horn (mid) spinal cord and was enriched in both excitatory (Excit-23, Excit-25, and Excit-30) and inhibitory (Inhib-15 and Inhib-21) clusters. Of note, several cluster “markers” of ventral cell types, such as Sim1, were not observed in adult spinal cord tissue and likely represent lingering RNA from developmental samples.

##### Excitatory (VE) (8 clusters)

This family of ventral, excitatory clusters was comprised of Excit-31, Excit-32, Excit-33, Excit-34, Excit-35, Excit-36, Excit-37, and Excit-38. Excit-31, Excit-32, Excit-33, and Excit-34 expressed low but positive levels of Lhx2, Lhx9, and Isl1, potentially suggesting dorsal dI1/dI2/dI3 embryonic lineages for these clusters. These clusters could be distinguished by Gm26673, Syt2, and Prlr (Excit-31), Mdga1 and Prlr (Excit-32), and Bnc2 and Pou6f2 (Excit-34). Excit-35, Excit-36, and Excit-37 are likely derived from the V2a lineage, as they expressed Vsx2 (Chx10) and included many cells from the Hayashi dataset that sorted cells based on Chx10 genetic expression. Excit-35 also expressed Vamp1, Pou3f1, Shox2, and Pou6f2 and Excit-36 expressed Esrrg. Intriguingly, many cells from the Baek dataset, which sorted cells based on spinocerebellar status were found in Excit-35, suggesting an important synaptic target of this population. Excit-37 expressed the V3 marker gene Sim1 as well as Rnf220. Supplemental Figure 3H.

##### Inhibitory (VI) (6 clusters)

This family of ventral, inhibitory clusters was comprised of Inhib-22, Inhib-23, Inhib-24, Inhib-25, Inhib-26, and Inhib-27. Each of these clusters expressed the glycinergic marker Slc76a5. Inhib-22 and Inhib-27 also expressed the gabaergic marker Gad2, Pax2, and Pou6f2. They were distinguished by low levels of Gata3 expression in Inhib-27, which may represent V2b lineage. Inhib-23 and Inhib-25 expressed Foxp2 and Esrrb, suggesting they correspond to the Foxp2 clade of V1 lineage neurons. They were distinguished by expression of Gm26673 and Pvalb in Inhib-23, which may suggest that this cluster included Ia-inhibitory neurons. Inhib-24 expressed both Pou6f2 and Nr5a2, suggesting that this cluster corresponded to the Pou6f2/Nr5a2 clade of V1 lineage neurons. Inhib-26 was the most robust ventral cluster and expressed the Renshaw marker genes Chrna2, Chrna7, and Calb1, suggesting that this cluster corresponded to Renshaw cells. Supplemental Figure 3I.

### Comparison to Two Previously Published Atlases

To determine how these neuronal clusters relate to previously characterized transcriptomic spinal cord cell types, we focused on the original clusters from the Sathyamurthy and Haring datasets because these two studies included a common set of cell types (dorsal horn neurons) and provided the most analysis, annotation, and marker validation for their respective cell types. First, we analyzed how cells/nuclei from the original studies were distributed into the new harmonized cluster of the meta-analysis (Figure 4A). Some ventral neurons from the Sathyamurthy dataset appeared in low-quality clusters that were discarded from the harmonized analysis due to low counts of genes per cell/nucleus and a lack of marker genes, whereas some neurons from the Haring dataset were classified as non-neural cell types or appeared in doublet clusters that were also discarded from the harmonized analysis. Nevertheless, we found that most original cell types fell within one of the harmonized neuronal atlas clusters or split into a small group of related neuronal clusters. The co-clustering between cells and nuclei from the original studies revealed many cell type similarities. For example, the majority of Haring Glut12 cells split into harmonized clusters Excit-14 and Excit-15, together with nuclei from Sathyamurthy DE-12 and DE-16 (Figure 4A). This is consistent with the original characterizations of these clusters, in that Haring Glut12 was principally marked by Grpr and Qrfpr, Sathyamurthy DE-12 was principally marked by Grpr, and Sathyamurthy DE-16 was principally marked by Col5a2 together with Qrfpr. In addition, prior comparison of the overall gene expression pattern of Haring Glut12 was most closely correlated with Sathyamurthy DE-12 and DE-16. This suggests that Glut12 and DE-12/DE-16 represent similar cell types that the Haring study kept as one cluster but which Sathyamurthy study split into two clusters. In the harmonized analysis and in the in situ hybridization validation above (Figure 3B,D), both Excit-14 and Excit-15 are relatively robust clusters (with robustness scores of 0.88 and 0.85, respectively) and can be distinguished by expression of Enpp1 in Excit-15, supporting the splitting of these related cell types into two distinct clusters.

**Figure 4.**
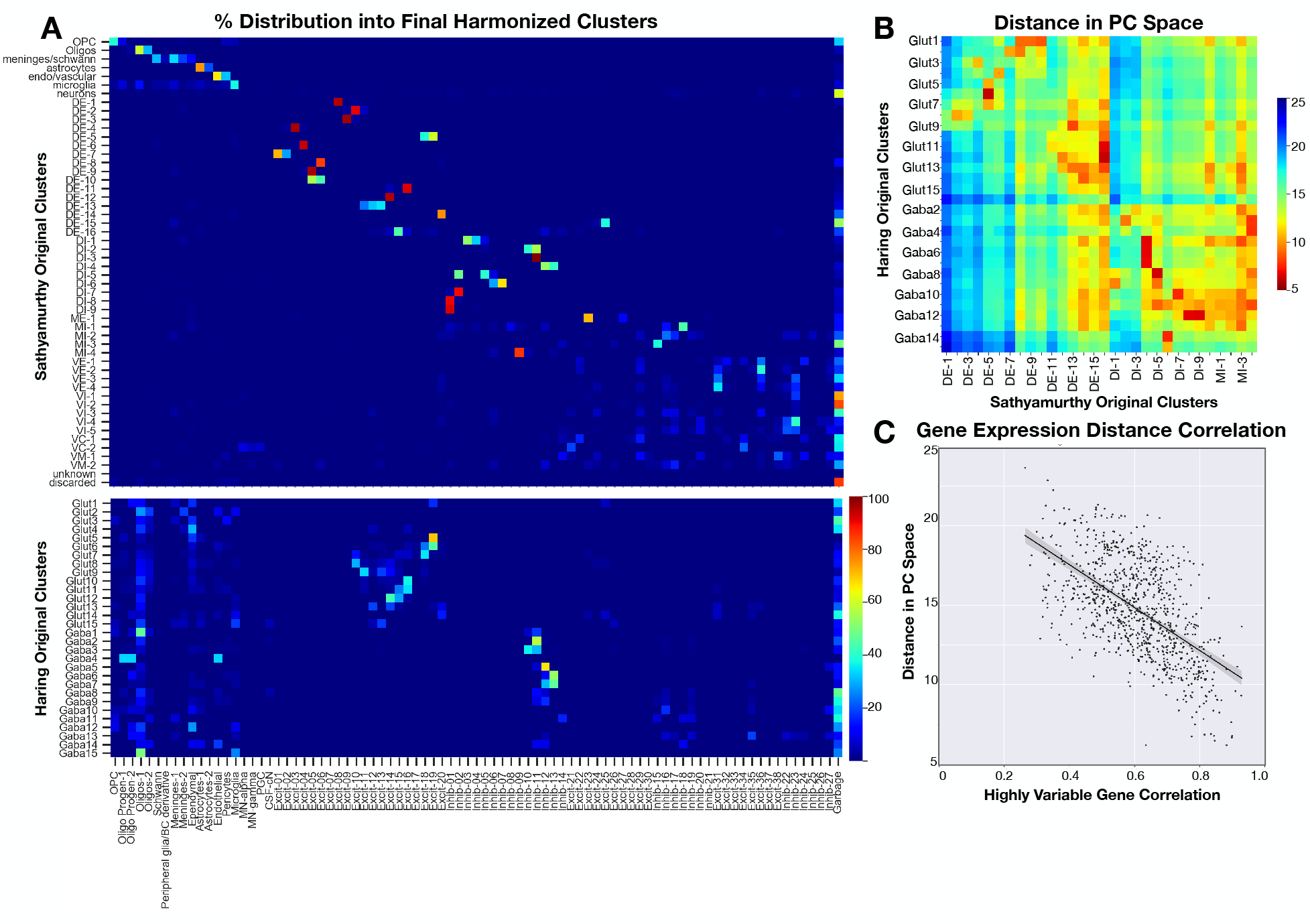
Relationship with two previously published spinal cord atlases. (A) The distribution of cells from the original clusters of the Sathyamurthy and Haring datasets (rows) into the harmonized clusters (columns), ranging from 0 blue to 100% red distribution. (B) The distance between the centroids of the cells/nuclei from the original Haring and Sathyamurthy clusters, measured in 50 dimensional principal component (PC) space. Only dorsal neuron clusters are shown for the Sathyamurthy dataset and in both datasets, every other cluster is labeled. Relatively short distances = red; long distances = blue. (C) Relationship between the distance in PC space and the correlation in gene expression between pairs of clusters from the Haring and Sathyamurthy datasets.

To compare the overall relationships between cells/nuclei from the Haring and Sathyamurthy studies with our harmonized meta-analysis, we calculated the distance in harmonized principal component “space” between the centroid of cells/nuclei from each original study’s cell types as well as correlation in expression of highly variable genes for each pair of cell types (Figure 4B,C). We found that increasing correlation between the original clusters’ gene expression strongly predicted closeness in the harmonized principal component space, suggesting that co-clustering in the harmonized analysis should accurately preserve and reveal relationships with the cell types described in the original studies (Figure 4C).

### Using Machine Learning to Classify Spinal Cord Cell Types

With this atlas of spinal cord cell types in hand, we next sought to establish a means to standardize and automate spinal cord cell type classification. First, we tested three strategies that have been used successfully to classify single cell data from other tissues on their ability to classify spinal cord cells into coarse cell types. These were label transfer^13^, a support vector machine, and a fully connected neural network (with two hidden layers of 512 nodes and L2 regularization for each). It is important to note that each of these models were trained using cell type labels from the harmonized analysis because there is no existing gold standard for spinal cord cell identities. In this context, the analysis that follows should be considered a feasibility study for machine learning classifiers on spinal cord single cell count data. The full merged dataset of 101,070 cells and nuclei was tested, including low quality cells and nuclei and doublets, in order to represent the full range of input raw data. All three strategies performed well, with label transfer showing the best performance (overall accuracy of 89%), followed by the neural network (83%), and then the SVM (80%) (Figure 5A and Supplemental Table 4).

**Figure 5.**
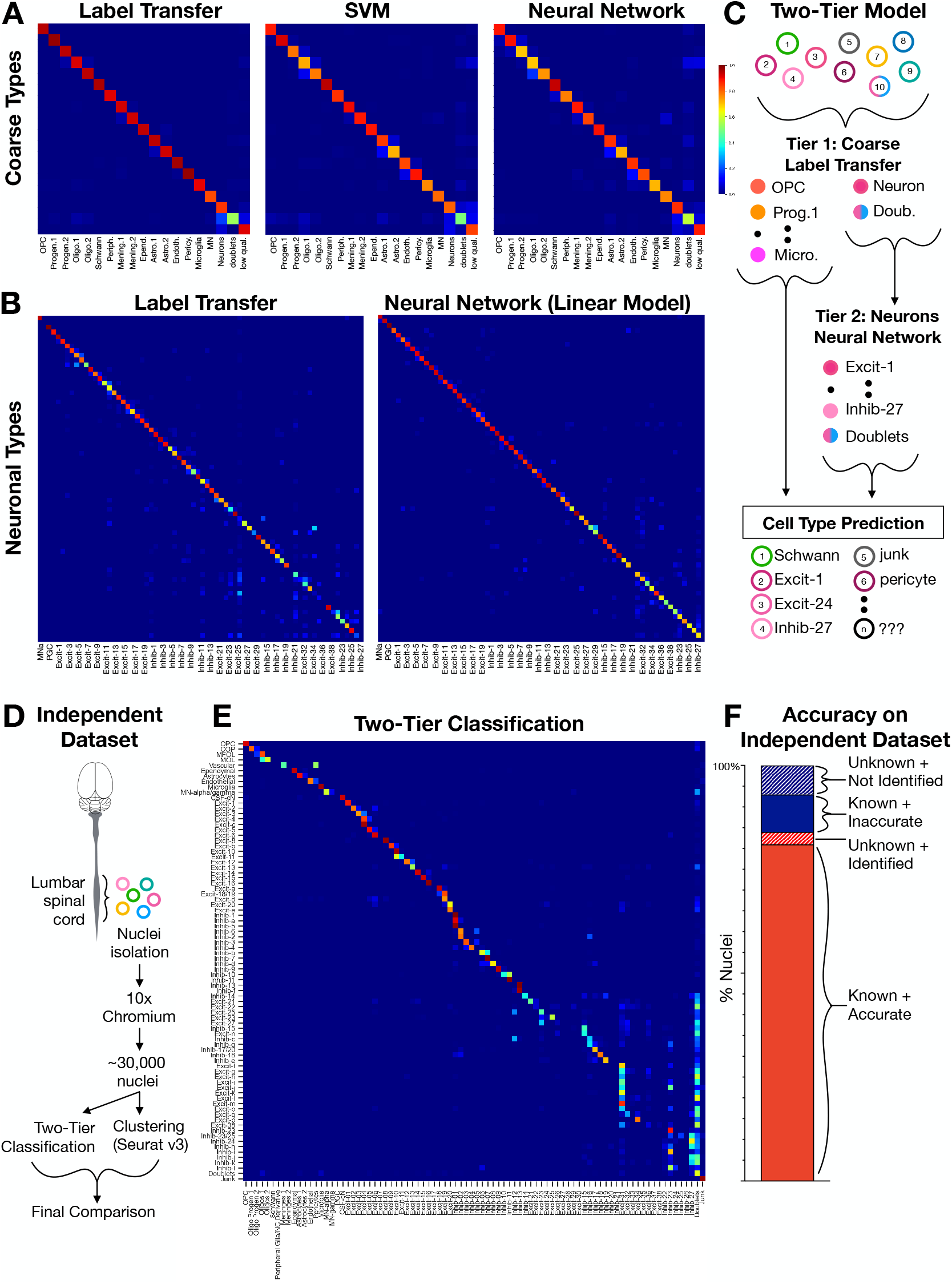
Computational classification of spinal cord cell types. (A) Confusion matrices of the F1 scores for the classification of coarse cell types using label transfer, a support vector machine (SVM), and a fully connected neural network (neural net), (blue = 0; maroon = 1). The actual cell types are in rows and the predicted cell types are in columns in the same order. (B) Confusion matrices of the F1 scores for the classification of fine neuronal sub-types using label transfer and a fully connected neural network. The actual cell types are in rows and the predicted cell types are in columns, both in the order presented in Table 1. Alternating cell types are labeled. (C) Model of the two-tiered classification approach in which all cells/nuclei are classified into coarse cell types using label transfer (also including low-quality “junk” and “doublets”). Subsequently, all cells/nuclei that were classified as neurons, motoneurons, or doublets by label transfer are further classified into 69 neuronal cell types (also including “doublets”). (D) Experimental design for generating an independent set of single nucleus RNA sequencing data. (E) Distribution plot showing how nuclei from each cluster (rows) were distributed into each of the harmonized cell types (columns), normalized by rows with dark blue = 0.0 fraction; maroon = 1.0 fraction). (F) Bar plot of the total counts of nuclei that were from “known” clusters and were correctly classified (81% of total), that were from “known” clusters and were incorrectly classified (9% of total), that were from “unknown” clusters but could be identified by their classification (3% of total), or that were from “unknown” clusters and could not be identified (7% of total). OPC=oligodendrocyte precursor cell; progen.1=oligodendrocyte progenitor 1; progen.2==oligodendrocyte progenitor 2; Olig.1=oligodendrocyte 1; Olig.2=oligodendrocyte 2; Periph.=peripheral glia; Mening.1=meninges 1; Mening.2=meninges 2; Epend.=Ependymal cells; Astro.1=astrocytes 1; Astro.2=astrocytes 2; Endoth=endothelial cells; Pericy.=pericytes; MN=motoneurons; low qual.=low quality. MNa=motoneurons alpha; PGC=preganglionic cell.

Next, we tested label transfer and neural networks on a more refined and challenging task: the classification of 69 neuronal sub-types. For label transfer, a two-tiered analysis was performed (dorsal sub-types and then mid/ventral sub-types) because we found that this approach was important for clustering spinal cord neurons. For the neural networks, a non-exhaustive handsweep of several hyperparameters was conducted, including network depth, optimizer, number of hidden nodes, and the number of training epochs and seven different models were tested (see Methods and Supplemental Table 4). We found that a linear model (with no regularization and with an SGD optimizer) showed the best performance, with an overall test accuracy of 85% (Figure 5B and Supplemental Table 4). The model showed very high confidence scores for correct predictions; however, performance varied with cell type prevalence suggesting a target for improving the model in the future (Supplemental Figure 4).

How should the performance of this model be viewed and should we expect automated classification to achieve 100% accuracy? Perfect performance would require perfect biological data: discrete cell types that express completely distinct patterns of gene expression and experimental data without doublets, low quality cells, or other sources of indeterminate data. Knowing that this is not possible, we still sought to determine a benchmark performance guide for the classification adult mouse spinal cord neurons using neural network models and considered four metrics of cluster definition and separation. We examined the relationship between the model performance for each cluster (F1 score) and (1) the co-clustering frequency of each cell type across 100 clustering iterations, (2) how distant each cluster was from its nearest neighbor in principal component space, and (3) the confidence with which clusters could be distinguished based on in situ marker expression (measured by in situ analysis sets of clusters) (Supplemental Figure 4). We found that the model performance varied with the co-clustering frequency of each cluster and with the ability to identify cell types in situ and we propose that these measures can be used to set a reasonable expectation for neural network performance. Overall, neuronal cells/nuclei of a given type co-clustered together 65% of the time (average from Supplemental Figure 2E) and a total of 70% of cells could be classified in situ (Supplemental Table 3). In comparison, the model’s accuracy of 85% reveals the outstanding performance of this approach.

To develop a standardized pipeline for classification of independent datasets unrelated to the original studies analyzed above, we considered a two-tiered approach that would take advantage of the strengths of both the label transfer for coarse classification (Tier 1) and a neural network model for classification of neuronal sub-types (Tier 2) (Figure 5C). We first selected all cells/nuclei that were assigned as doublets or neurons during the harmonized analysis above to represent the output of the first-tier and input to the second tier. In this context, we trained another set of five neural network models (see Methods and Supplemental Table 4). A neural network model with one hidden layer (256 nodes) and SGD optimizer showed the best performance (overall accuracy of 80%) and was selected for further work.

As a final performance test of the two-tiered model, we applied it to spinal cord nuclei from an independent experiment. Nuclei were isolated from the lumbar spinal cords of four adult mice, sequenced using 10x Chromium, clustered using Seurat, and marker genes were identified for each cluster (Figure 5D). 90% of nuclei (out of 28,584 total) were in clusters that could be assigned a cell-type label based on user-based marker gene expression (“known” clusters). In cases for which labels could not be confidently assigned (10% of nuclei, “unknown” clusters), a placeholder name was given. We performed classification of all nuclei from the independent dataset that passed quality-control thresholds (Figure 5C) in an analysis that took less than thirty minutes of computational time (∼20 minutes for Tier 1 and less than one minute for Tier 2).

We found that 90% of nuclei from “known” clusters were accurately classified by the two-tiered model (Figure 5F “known + accurate”). We next considered how this model performed upon the classification of nuclei from the challenging “unknown” clusters that could not be identified based on marker genes. Surprisingly, we found that 28% of unknown nuclei could be identified with the two-tier classification model (Figure 5F “unknown + identified). Thus, the two-tiered model surpassed the ability of experienced users to identify spinal cord cell types.

Of note, several cell types were not expected to be present in the independent dataset, including Schwann cells, peripheral glia and meninges 2 (based on the surgical dissection method used that did not include spinal roots or outer layers of meninges) and including PGC, Excitatory-7, and Excitatory-17 (based on the lumbar region and adult age that was used). As expected, these cell types were not predicted by the two-tiered model. There were also several cell types that were not classified as expected. In particular, several mid/ventral cell types were not detected in the independent dataset while two ventral clusters (Excitatory-31 and Inhibitory-27) were over-represented. This may reflect a training dataset that is not large enough to train a model that distinguishes closely related cell types, that small cell types are not modeled as well, and that some mid/ventral clusters are defined partly by early postnatal gene expression contained within the harmonized analysis but absent from the independent adult dataset.

These results establish a two-tiered model based on label transfer and a neural network as an effective approach for the computational classification of single cell sequencing data, even in the context of the finely separated populations of spinal cord neurons. The neural network model was at least as accurate as other methods such as Seurat-based clustering and high-content in situ hybridization and was orders of magnitude faster. In addition, it can standardize spinal cord cell type classification so that a unified and harmonized set of cell types can be identified and studied consistently between datasets, biological conditions, and laboratories throughout the field.

### SeqSeek: A Community Resource for Analyzing and Classifying Spinal Cord Cell Types

Finally, we have developed an online resource for spinal cord single cell data, SeqSeek (available at seqseek.ninds.nih.gov). This resource includes user-friendly tools to search gene expression across spinal cord cell types using single genes or gene lists and to view spatial distributions of selected marker genes (SeqSeek Genes), to compare gene expression between clusters or groups of clusters (SeqSeek Cell-Types), and to access the SeqSeek algorithm for cell type classification (SeqSeek Classify).

## DISCUSSION

For the field of spinal cord biology to build upon the incredible promise of single cell technologies, it is critical to establish a standard set of cell types. Here, we leveraged and expanded upon the previously published single cell sequencing studies of the postnatal mouse spinal cord to define 84 types of spinal cord cells. We present a harmonized atlas of these cell types; a validated combinatorial panel of markers to facilitate their study either *in vivo*, in tissue sections, and *in vitro* cell culture; computational resources for classifying spinal cord cells based on transcriptomics; and a web-based resource, SeqSeek, to allow the community to interact with and explore single cell spinal cord data. This work establishes a common framework that will serve as a powerful resource for the field and facilitates the discovery of new biological features of spinal cord cell types.

The first key consideration for this atlas is whether the cell types of the atlas are correct. In the absence of a commonly accepted standard set of spinal cord cell types, it is impossible to answer this question completely. However, several pieces of evidence support the accuracy of the harmonized clusters. First, these clusters are robust to different clustering approaches, suggesting that they reflect underlying biological signatures rather than a technical artifact. Second, these clusters correspond well with prior gene expression studies of the postnatal spinal cord, including three single nucleus sequencing datasets that were not included in the harmonized clustering: an independent dataset that we clustered separately and used to test the SeqSeek Classify algorithm, and two very recent studies that found similar markers to the harmonized set^8,9^. Third, and most importantly, nearly all of the predicted marker neuronal co-expression patterns could be validated in tissue and several represent well-established molecular markers of accepted “cell types”.

In addition to serving as a powerful reference resource, what new biological information can this study reveal? By incorporating the analysis of six independent studies we have been able to resolve cell types at a granular level and created the most comprehensive description of spinal cord cell types to date. In particular, the increased power from studying many neurons across postnatal development allowed us to better characterize mid and ventral cell types. While these clusters still display low to moderate robustness, this is mainly because they are highly related to each other through overlapping gene expression patterns. Previously, we noted this trend amongst ventral clusters and we now identify spatial patterns of gene expression (such as Pou6f2 and Nfib) as a source of this relatedness. We propose that the combination of embryonic lineage and settling location contribute to the definition of cell types in the mid and ventral horn regions. This in turn gives rise to both cell type heterogeneity and the overall similarity of the mid area and ventral horn clusters.

Another type of new biological insight is based on the co-clustering of cells defined by different parameters. For example, the largest fraction of neurons from Hayashi et al., which isolated V2a lineage derived neurons co-clustered within Excit-35 together with the largest fraction of neurons from Baek et al., which isolated spinocerebellar neurons. This co-clustering suggested that these cells are highly similar and may link V2 embryonic origin with spinocerebellar circuit connectivity. In support of this connection, the established V2a marker genes Shox2 and Sox14 were both identified as markers of putative lamina VII spinocerebellar tract neurons in the original Baek et al. study. Thus, co-clustering of cells across different studies can reveal candidate linkages across cell type features and illustrates the power of a harmonized atlas across time and biological conditions.

This study also highlights important experimental and analytical parameters. On the experimental side, this study revealed the differences between using cells versus nuclei for transcriptomic profiling. As expected, we found that single cell studies detected more genes per cell than single nucleus studies did per nucleus, but that single cells also showed higher levels of stress response gene expression. Unexpectedly, we also found that the major single cell atlas of the juvenile mouse nervous system failed to include any ventral interneurons or motoneurons while these were found readily even in adult tissue that used single nuclei. Whether this reflects greater vulnerability of ventral cells to tissue dissociation and cell stress, or whether other technical limitations were present, remains to be determined.

On the analytical side, this work is among the first practical applications of automated classification for large and complex single cell datasets. A wide range of cell annotation approaches have been described recently but it is not yet clear which methods will work best for each type of data^14-18^. A comparative analysis of automated classification approaches across diverse datasets found that SVM and neural network models showed the best performance on the Allen Brain Atlas dataset of 92 neuronal cell types – a dataset similar in scale and complexity to the harmonized analysis here^18^. This analysis also found that performance depends partly on the number of cell types and the “complexity” (the relatedness between clusters) of a dataset, similar to what we observed.

The described here displayed excellent performance in the computationally challenging task of classifying cells and nuclei into the 69 “fine” resolution neuronal cell types of the spinal cord. In the future, larger spinal cord single cell datasets will be available and the neural network model that we presented here can be refined and improved. Specifically, larger training datasets may facilitate classification of closely related mid/ventral neuronal populations; region or sample age specific training datasets may reduce the number of cell types that cannot be detected; and generative models may be used to enhance training on rare cell populations. As this work proceeds, we expect that increasingly powerful neural network models will be developed that allow rapid, accurate, and standardized classification of all spinal cord cell types directly from raw sequencing data. This could be done by individual users with downloadable models or through the development of a spinal cord single cell data commons that could continuously refine the models and provide classification analysis through a cloud-based platform, similar to what has been proposed for the Human Cell Atlas^19^. A forthcoming study aims to partially address these challenges. Theis and colleagues propose a method called *single-cell architectural surgery* that uses transfer learning to map query datasets onto a reference, simultaneously contextualizing the query while updating the reference. This allows for decentralized reference building without the sharing of raw data, which could further increase effectiveness of neural network-based classifiers^20^.

There are several notable limitations to this study and to single cell transcriptomics in general. Most specifically, this analysis is limited in scope to RNA expression in the postnatal mouse spinal cord. As more data become available from studies that include more specific regions of the spinal cord, more biological conditions, more developmental stages, more species, more specific cellular features, and more -omics modalities, we anticipate that this work will reveal exciting new insights from single cell data. As examples, future work could incorporate embryonic single cell data^7^ and lineage tracing to link together developmental origin with postnatal cell types or could focus deeply on specific spinal cord regions and cell types. Indeed, forthcoming work has revealed an impressive diversity of PGC visceral motoneurons that are enriched in either the thoracic or sacral spinal segments^21,22^. Relatedly, the in situ hybridization experiments here are also limited in scope, being specific to the adult lumbar spinal cord. The failure to detect several genes could reflect that these genes are no longer expressed at the adult stage or lumbar region that we analyzed, that the cell types themselves are not present (being transiently found in early postnatal stages or only in other spinal cord regions), or technical issues. As new data and technologies become available, we anticipate an explosion of single cell data and the opportunity to periodically supplement, evolve, revise, and refine the work presented here.

A second notable caveat is that this analysis is all population based. Data is captured from thousands of individual cells, but the rate of false negative data in each cell and the requirement for statistical power necessitates analyzing many cells of each type and considering population level shared patterns. It is likely that by emphasizing common patterns, this analysis underrepresents true biological variability, including “noisy” gene expression and continua of cell types. For example, three very different methods – single cell data clustering, multi-plexed in situ hybridization, and an artificial intelligence neural network – all showed a relatively weak ability to classify ventral cell types into discrete types and a relatively strong but still imperfect ability to classify dorsal cell types. We propose that this reflects some technical limitations but also a fundamental complexity and diversity in how gene expression is controlled within individual cells and in cell type populations.

Finally, it is crucial to note that single cell/nucleus profiling, particularly single cell/nucleus RNA sequencing, produces one perspective on cell types and it is not yet clear how this will relate to other core cellular features such as developmental lineage, circuit connectivity, electrophysiology, and behavioral function. Re-considering the very definition of “cell type” and identifying the most useful system for classifying cells is now a fundamental task in understanding nervous system function. We expect that in each tissue, indeed in each region of each tissue, there may be different organizing principles of “cell types”. In that context, the work here provides a comprehensive atlas of spinal cord transcriptomic cell types that can be used as a framework to compare with other cellular features.

Overall, this work brings together the first six single cell studies of the post-natal mouse spinal cord to create a standard reference set of spinal cord cell types. It will (1) serve as a unifying resource and nomenclature for the field, (2) provide a validated and combinatorial set of markers that can be used to translate this rich sequencing data back into tissue based studies, (3) be a template for the computational analysis of single cell data from complex neural tissue, and (4) facilitate the community-wide use of single cell data through a web-based resource. We hope that this work will facilitate the design and interpretation of cell-based studies of behavior and will open up opportunities for many new discoveries.

## METHODS

### Mice

Animal experiments were performed in accordance with institutional guidelines and approved (protocol #1384) by the National Institute of Neurological Disorder and Stroke’s Institutional Animal Care and Use Committee. An even balance of male and female mice that were 9 weeks old and of mixed C57BL/6J and BALB/cJ background were used for single nucleus sequencing (four mice) and validation studies (six mice).

### Published Data Acquisition

Published data were downloaded from the NCBI Sequence Read Archive (SRA). Raw datasets were used instead of investigator-provided count matrices so that we could align all sequences to the same genome and apply uniform data filtering. All raw datasets were pre-processed using technique-specific pipelines. For data from Sathyamurthy et al. (DropSeq, GEO:GSE103892, SRA:SRP117727), data were downloaded in fastq format from SRA. A count matrix was created following the steps in the McCarroll lab DropSeq cookbook^23^. For data from Hayashi et al. (GEO: GSE98664, SRA: SRP106644) and Zeisel et al. (SRA: SRP135960) both 10X, 10X sequence data were download from SRA in BAM format then converted to cellranger-compatible fastq files using the 10X-provided bamtofastq tool^24^. Count matrices were created using the 10X cellranger count tool^25^. Data from Haring et al. (C1 Fluidigm, GEO: GSE103840, SRA: SRP117627) were downloaded from SRA. Each cell had its own fastq file for a total of 1545 files. We followed the UMI tools single cell tutorial^26^ to remove the UMI and process the sequences. For the Rosenberg et al. data (SplitSeq, GEO: GSE10823, SRA: SRP133097), data were downloaded in fastq format. Count matrices were made using the split-seq-pipeline tool developed by the Seelig Lab^27^. The STAR alignment tool within cellranger (v020201) was used to align the sequences from each dataset to a reference genome that was custom built to include all introns and exons.

### Merged Analysis and Clustering

Count matrices for each dataset were merged to obtain the full data file and we then applied uniform data filtering across the merged file. We analyzed all cells and nuclei with at least 200 detected genes (to exclude low quality or “empty” barcodes) and with less than 5% of transcripts being mitochondrial (to exclude lysing cells or mitochondria-nuclei doublets). This yielded over one hundred thousand total cells/nuclei. Of note, by starting with the raw data and setting relatively relaxed thresholds for data inclusion, we analyzed more cells/nuclei from several of the original studies than were analyzed in the corresponding published datasets. The merged data was analyzed using Seurat v3. Clustering was performed in three phases on (1) all cell types, (2) all neurons, (3a) presumptive ventral neurons and (3b) motorneurons. For phase 1, data integration was performed by study, 2,000 highly variable genes were detected, and the most significant principal components were identified by elbow plot and manual inspection of the contributing gene lists and 28 PCs were used for clustering. To select cluster resolution, a range of values were tested from 0.2 to 8 and cluster evolution or clustree plots were used to determine when cluster splitting stabilized, and resolution 1.2 was selected. For phase 2, raw data from all cells in neuronal clusters was used, re-scaled, re-normalized, and re-integrated, the top 4,000 highly variable genes were detected and the top 40 PCs were selected (using the approach described above). Resolutions from 0.8 through 10 were tested and a resolution of 8 was selected. A third phase of targeted sub-clustering was done because mid/ventral and motoneuron sub-types did not separate well in preliminary neuron analysis. Indeed, the robustness scores for mid/ventral cell types were very low until they are analyzed in a focused principal component space (Supplemental Figure 2). For phase 3a, presumptive ventral neurons were identified by markers and by coalescence on uMAP into a central “blob” and for phase 3b, motorneurons were identified by expression of classic markers (Chat, Isl1, Prph). In each case, the procedures described above were used to sub-divide these cell types and the following parameters were used: 3a: 40 PCs, resolution 4; 3b 7 PCs, resolution 0.6.

For all three phases, each cluster was analyzed for candidate marker genes and excluded if the cluster met either of the following criteria. Clusters were considered “low-quality” if they had fewer than three significant markers relevant to cell type, particularly if they showed very low nGene. Clusters were considered “doublets” if they had significant markers for multiple unrelated cell types and a “barnyard” plot of the top ten markers of each cell type showed that individual cells in the cluster displayed both sets of markers. For all three phases, we used the following method to determine whether candidate pairs of clusters should be merged: a dendogram based on mean gene expression and UMAP location were used to systemically identify closely related clusters and we then probed for differential gene expression. Pairs with fewer than three genes enriched in each cluster (six total) were merged unless a “classic” marker gene from the literature was one of five differentially expressed genes. Cell type annotations for the non-neuronal cell types were based on the presence of well-established marker genes (Supplemental Table 1) and on the gene expression patterns in the Allen in situ hybridization database (for meningeal, ependymal, Schwann cell and peripheral glia clusters).

The meta-data (and associated final cell labels) are available in Supplemental Table 5.

### Cell Type Relationships and Comparison with Prior Studies

To examine the relationship between the 69 neuronal clusters in the harmonized analysis, the centroid of each cluster was calculated by grouping the cells by their labels and determining the mean of each PC. Then, the pairwise Euclidean distance between each cluster was calculated using 50 PCs. This was passed to the stats::hclust function using method = “complete”. The final dendrogram was plotted using the graphics::plot function.

To examine the distribution of the original Haring and Sathyamurthy clusters amongst the harmonized clusters, the frequency of each pair-wise combination of original and harmonized clusters was counted. These data were then pivoted to wide form to produce the matrix with harmonized clusters along the x-axis and original clusters along the y-axis. Finally, the data was row-normalized, so that the color represents the fraction of the original label occurring in each harmonized cluster.

To examine the distance between the original Haring and Sathyamurthy clusters in harmonized PC space, the pairwise distance between the centroids of the original clusters was calculated as above. Small distances, representing close clusters, are displayed with hot colors, while large distances, representing far apart clusters, are displayed with cold colors.

To examine the correlation between PC distance and the expression of the 500 most highly variable genes in the harmonized data, the average expression of these genes was calculated for each original cluster, which yielded two matrices: one a genes by cluster matrix of the Haring data, and the other a gene by cluster matrix of the Sathyamurthy data. The correlation of gene expression in each cluster between these matrices was calculated using the lineup::corbetw2mat function (CRAN version 0.37.11). These correlation scores were then plotted against the PC distances calculated above. A linear regression with 95% confidence intervals is shown.

### RNA In situ Hybridization

14 µm fresh frozen spinal cord sections from segment L4 on Leica Apex slides were used with a set of 97 RNAScope HiPlex probes (Supplemental Table 2) from ACDBio, according to the manufacturer’s instructions. Images for each set were registered using RNAscope HiPlex Image Registration Software and brightness/contrast were adjusted using Adobe Photoshop. Counting of cells for each set were done as follows. Set 1: All Chat+ cells in any laminae. Set 2: Any dorsal cell that expressed any of Cpne4, Maf, or Prkcg. Set 3: Any cell in the dorsal horn with any of Slc17a6, Rreb1, Reln, or Car12. In addition, Gbx2 cells were counted separately amongst any cell in the deep dorsal horn with Slc17a6. Set 4: Any cell in the dorsal horn with any of Col5a2, Enpp1, Sox5, Tac1, Tac2, Nmu, Megf11, Mdga1, Pmfbp1, or Onecut2. Set 5: Any cell in laminae 1-4 with any of Slc6a1, Gad2, or Kcnip2. Set 6: Any cell in the dorsal horn with any of Mlxipl, Pdyn, Gal, Npy, Qrfpr, Sstr2, or Rspo3. Set 7: Any cell in laminae 4-6 with any of Slc17a6, Adamts2, Lmx1b. Set 8: Any cell in laminae 4-6 with either Slc6a5 or Gad2. Set 9: Any cell in laminae 6-8 with Slc17a6. Set 10: Any cell in laminae 6-8 with any of Pax2, Slc6a5 or Gad2. The number of cells counted in each set are listed in Supplemental Table 2 and were from one section per animal, though multiple sections per animal were inspected for expression pattern consistency. Sections from three animals (2 male and 1 female or 2 female and 1 male) were counted for each set.

### Single Nucleus Sequencing

Nuclei were obtained as previously described^28^ and were processed for single cell sequencing using the 10X Genomics Chromium Single Cell 3’ Kit (v3 chemistry) and sequenced at a depth of approximately 50,000 reads per nucleus. Clustering was performed as described above and cluster identities were determined using the combinatorial marker code in Table 1 where possible (“known clusters”). Clusters that could not be identified in this manner were analyzed for neurotransmitter status and given a placeholder identification (“unknown clusters”).

### Computational Classification

#### Label Transfer

Label transfer analysis was performed using Seurat v3(.1.5). For both coarse cell types and clean neurons, 10% of cells were withheld as the query dataset, whilst the remaining were used as the reference dataset. Broadly, label transfer consists of two-steps. First, the transfer anchors are identified using the FindTransferAnchors function. Second, these anchors are then used to transfer cluster labels to the query dataset with the TransferData function.

For label transfer of coarse cell types, FindTransferAnchors was called with reduction = “pcaproject”, dims = 1:28, and npcs = NULL to project the previously calculated PCA onto the query data using the same dimensions as were used in clustering the reference data. TransferData was also called with dims = 1:28 for the same reason.

Label transfer of clean neurons was performed in a two-step process. First, all cells in mid- or ventral-clusters were grouped as one cluster. Then, the dorsal-clusters were transferred along with one “mid/ventral” cluster. Second, those cells classified as “mid/ventral” were labelled using only neurons from mid- or ventral-neuron clusters. In each case, a new reference object was created from the appropriate cells – all neurons for step 1 and mid-/ventral-neurons only for step 2 – via integration, as previously discussed in “Merged Analysis and Clustering”. Label transfer was run as described for coarse cell types, with the exception that dims = 1:100 was set for all neurons, and dims = 1:30 was set for mid-/ventral-neurons.

In the final two-tier analysis, label transfer was performed as discussed for coarse cell types. Any cells labelled “Neuron”, “Motorneuron”, or “Doublets” were passed to the neural network for further classification. The decision to include doublets for further classification was founded on the observation that a non-trivial number of neurons were mis-classified as doublets at the coarse cell-type level.

#### Support Vectror Machine

Support vector machine analysis was performed using scikit-learn version 0.22.2.post1. Count matrices were taken from the default Seurat RNA assay count slot as sparse matrices. Cluster labels were numerical encoded with LabelEncoder(). To preserve sparsity for reduced training time, these counts were scaled with MaxAbsScaler(copy=False). As LinearSVC() is known to be a faster and more scalable than SVM(kernel=“linear”), it was selected for use^29^. As the number of samples was significantly greater than the number of features, the dual parameter was set to “False”^30^. Finally, to help ensure convergence, the max_iter parameter was increased from the default of 1000 to 10000. This pipeline achieved an overall accuracy of 80% on the validation data. Though this performance could likely be improved by hyperparameter tuning, given the performance of alternative models, the support vector machine was not selected for further use.

#### Neural Networks

Count Matrices were taken out of the default Seurat RNA assay count slot as sparse matrices. The counts were log x+1 transformed then scaled by the maximum number of counts for any gene in a cell. The data were converted into TensorFlow sparse tensors for input into neural networks define via the Keras interface to TensorFlow. Hyperparameters were initially set to default values, with a network structure consisting of direct connections between the input and output nodes. This simple linear model was the baseline. We added additional layers from 1 to 4 hidden layers, at various widths from 16 nodes to 512 nodes in a layer. The optimizer we switch from the default “Adam” optimizer to singular gradient descent (sgd). L1, L2 and dropout regularization were attempted. Additionally, various batch sizes were tested. Initially, networks trained for coarse analysis used a batch size of 128 to speed training. Whereas the training was faster, validation accuracy improved by around 5% when we lowered the batch size to 32. No additional improvement was seen at a batch size of 16, so the batch size was set to 32 for the rest of the study. In general, we used the learning curves to guide the changing of hyperparameters^31^.

For the analysis of coarse cell types (Figure 5A), a model with two hidden layers of 512 nodes each and L2 regularization was used. For the analysis of the neuronal sub-types (Figure 5B), seven models were tested: (1.1) a linear model with no regularization (1.2) a linear model with L2 regularization (learning rate 0.001) (1.3) a neural network with two hidden layers of 512 nodes each (1.4) an ensemble-like neural network with one hidden layer (128 nodes and L2 regularization) and two hidden layers that were concatenated, (1.5) a neural network model with three hidden layers (512, 256, 128 and L2 regularization on the 512 node hidden layer (1.6) a neural network model with 3 layers (128, 128, 128 and L2 regularization on the first hidden layer) and (1.7) a linear model with no regularization with an SGD optimizer. Interestingly, the baseline model had the largest validation accuracy. Since the training accuracy is 100% as compared to 85% in the validation set, the model is clearly over fitting the training data. Adding regularization helped to lower the gap between the training and validation accuracy, but the overall validation and test accuracies are still lower suggesting that the over trained model will perform better on unseen data. Additional work to improve this model is needed and adding more data from new experimental studies in the future will help improve the validation accuracy. For the analysis and training of neurons and doublets together (Tier 2), five models were tested: (2.1) a linear model with no regularization (2.2) a linear model with L2 regularization (2.3) a neural network model with one hidden layer of 128 nodes (2.4) a neural network model with one hidden layer of 128 nodes and SGD optimizer, and (2.5) a neural network model with one hidden layer of 256 nodes and SGD optimizer. The final model (2.5) was selected for Tier 2.

In the analysis of “unknown clusters” (Figure 5F), individual nuclei were “identified” if (1) they were from an “unknown” cluster and were classified into a harmonized true cell type (not “junk” or “doublets”) and (2) at least 80% of the total nuclei from their cluster of origin were classified into the same single harmonized cell type.

## Data Availability

Raw sequencing data from single nucleus sequencing will be available for download at GEO upon publication. A searchable version of all data is available www.seqseek.ninds.nih.gov and links to all raw data will be available at the same site. Associated code is available at https://github.com/ArielLevineLabNINDS.

### Acknowledgements

We are grateful to Dr. Vilas Menon (Columbia University) for his advice throughout this project and to Mr. Stefan Stoica for technical assistance with the validation data analysis. This work was supported by the Intramural Research Program of the NIH, NINDS and CIT. This research was supported in part by an appointment to the National Institute of Neurological Disorders and Stroke Research Participation Program administered by the Oak Ridge Institute for Science and Education (ORISE) through an interagency agreement between the U.S. Department of Energy (DOE) and the National Institute of Health. ORISE is managed by ORAU under DOE contract number DE-SC0014664. All opinions expressed in this paper are the author’s and do not necessarily reflect the policies and views of NIH, NINDS, DOE, or ORAU/ORISE.

## Author contributions

D.E.R, K.J.E.M, and A.J.L conceived of this project. D.E.R, S.C.K., and A.J.L carried out the merged analysis and comparison of cell types with the literature. D.E.R. and R.B.P.C. carried out the cell type analysis, study comparison analysis, and algorithm design and testing. L.L. carried out the in situ hybridization experiments. R.B.P. C. and A.J.L wrote the manuscript and prepared figures, with help from D.E.R., K.J.E.M., and S.C.K. All authors contributed to editing the final manuscript.

