## Supplemental Figures for "A Harmonized Atlas of Spinal Cord Cell Types and Their Computational Classification"

### Supplemental Figure 1 - related to Figure 1

**A**

| Study Order | Reference | Age | Mouse Strain | Tissue | Cells or Nuclei | Barcoding Method |
|---|---|---|---|---|---|---|
| 1 | Sathyamurthy et al., 2018 | 8 weeks | Mixed C57BL/6 and BALB/c | full tissue lumbar | nuclei | DropSeq |
| 2 | Hayashi et al., 2018 | perinatal | CB6F1/J | sorted Chx10+ cells cervical and lumbar | cells | 10X Chromium |
| 3 | Haring et al., 2018 | 3-4 weeks | C57BL6/N and C57BL6/J | sorted dorsal horn neurons region not specified | cells | C1-SQRT |
| 4 | Rosenberg et al., 2018 | 2 and 11 days | C57BL/6 and DBA/2 | full tissue region not specified | nuclei | SplitSeq |
| 5 | Zeisel et al., 2018 | 3 weeks | CD1 and Swiss Webster | full tissue cervical to sacral | cells | 10X Chromium |
| 6 | Baek et al., 2019 | 1 week | FVB | sorted spinocerebellar cells cervical and thoracic | cells | Single Well |

**B**

| | Sathyamurthy | Hayashi | Haring | Rosenberg | Zeisel | Baek |
|---|---|---|---|---|---|---|
| **OPC** | 322 | 3 | 8 | 950 | 790 | |
| **Oligo Progen-1** | 59 | 2 | 10 | 706 | 582 | 1 |
| **Oligo Progen-2** | 167 | | 25 | 469 | 629 | |
| **Oligos-1** | 3822 | 5 | 118 | 943 | 2832 | 3 |
| **Oligos-2** | 807 | | 30 | 470 | 922 | |
| **Schwann** | 1165 | | | 9 | 1 | |
| **Peripheral glia/ NC derivative** | 298 | | | 2 | 1 | |
| **Meninges-1** | 1661 | 5 | 6 | 444 | 18 | |
| **Meninges-2** | 641 | | 6 | 111 | | |
| **Ependymal** | 468 | 2 | 52 | 260 | 249 | |
| **Astrocytes-1** | 2219 | 6 | 13 | 1457 | 1907 | 2 |
| **Astrocytes-2** | 443 | 3 | 1 | 694 | 309 | |
| **Endothelial** | 1075 | 3 | 15 | 393 | 1462 | |
| **Pericytes** | 463 | | 9 | 310 | 826 | |
| **Microglia** | 466 | 9 | 17 | 438 | 640 | 16 |
| **Neurons** | 5473 | 421 | 1042 | 11201 | 1130 | 86 |
| **TOTAL** | **19549** | **459** | **1352** | **18857** | **12298** | **108** |

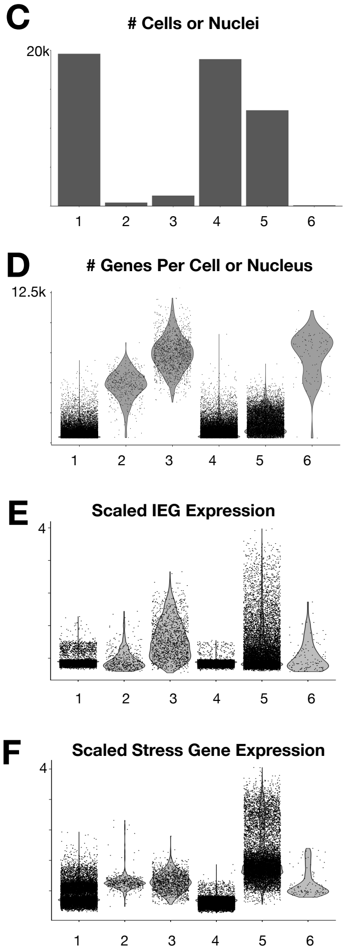


**Supplemental Figure 1 (related to Figure 1). Integrated analysis of six independent studies to reveal spinal cord coarse cell types.** (A) Table presenting the six studies that were merged for the harmonized analysis, arranged in publication order. (B) Table presenting the number of cells/nuclei from each study that were present in each of the coarse cell type clusters. Low-quality nuclei from the Sathyamurthy dataset are included, though they were discarded as such or labeled as neurons in preliminary analysis but discarded later in the neuron sub-analysis. (C) The cells/nuclei from each study varied in terms of the total numbers included in the final set of harmonized clusters (# cells/nuclei, scale is 20,000), the number of genes detected per cell/nucleus (scale is 12,500), and the expression of immediate early genes (IEG) and stress related genes (scale is 4 for both and the data are presented as the scaled expression of the gene module normalized to 100 randomly selected genes).

### Supplemental Figure 2 - related to Figure 2

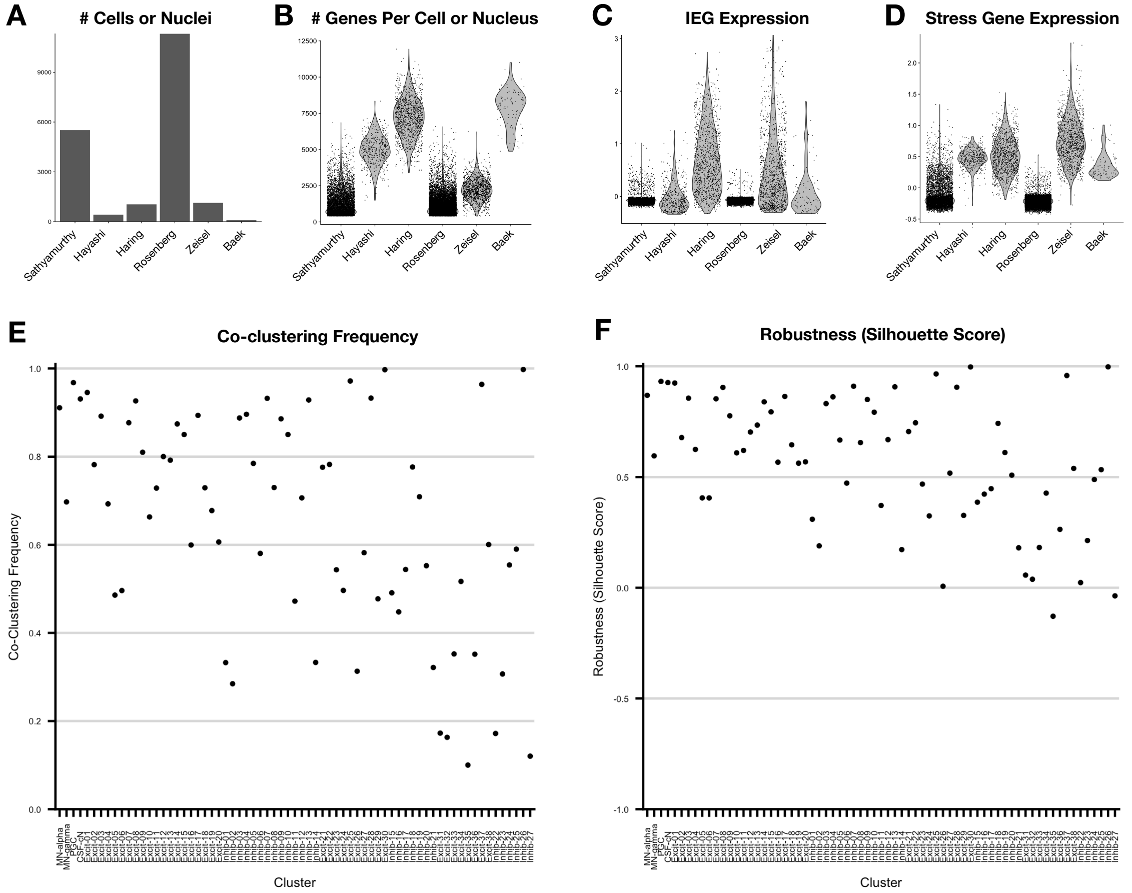


**Supplemental Figure 2 (related to Figure 2). Integrated analysis of six independent studies to define 69 spinal cord neuron cell types of varying robustness.** (A-D) The cells/nuclei from each study varied in terms of the total numbers included in the final set of 69 neuronal clusters (# cells/nuclei), the number of genes detected per cell/nucleus, and the expression of immediate early genes (IEG) and stress related genes (scale is 4 for both and the data are presented as the scaled expression of the gene module normalized to 100 randomly selected genes). The studies are presented in publication order. (E) The co-clustering frequency of the cells/nuclei from each cluster when clustering was automated and run 100 times using a random 80% of the dataset each time, analyzed in two tiers (first tier: mid/ventral grouped together; second tier: mid/ventral sub-types only). (F) The "robustness score" (the silhouette value of the co-clustering frequency matrix) of each cluster is shown.

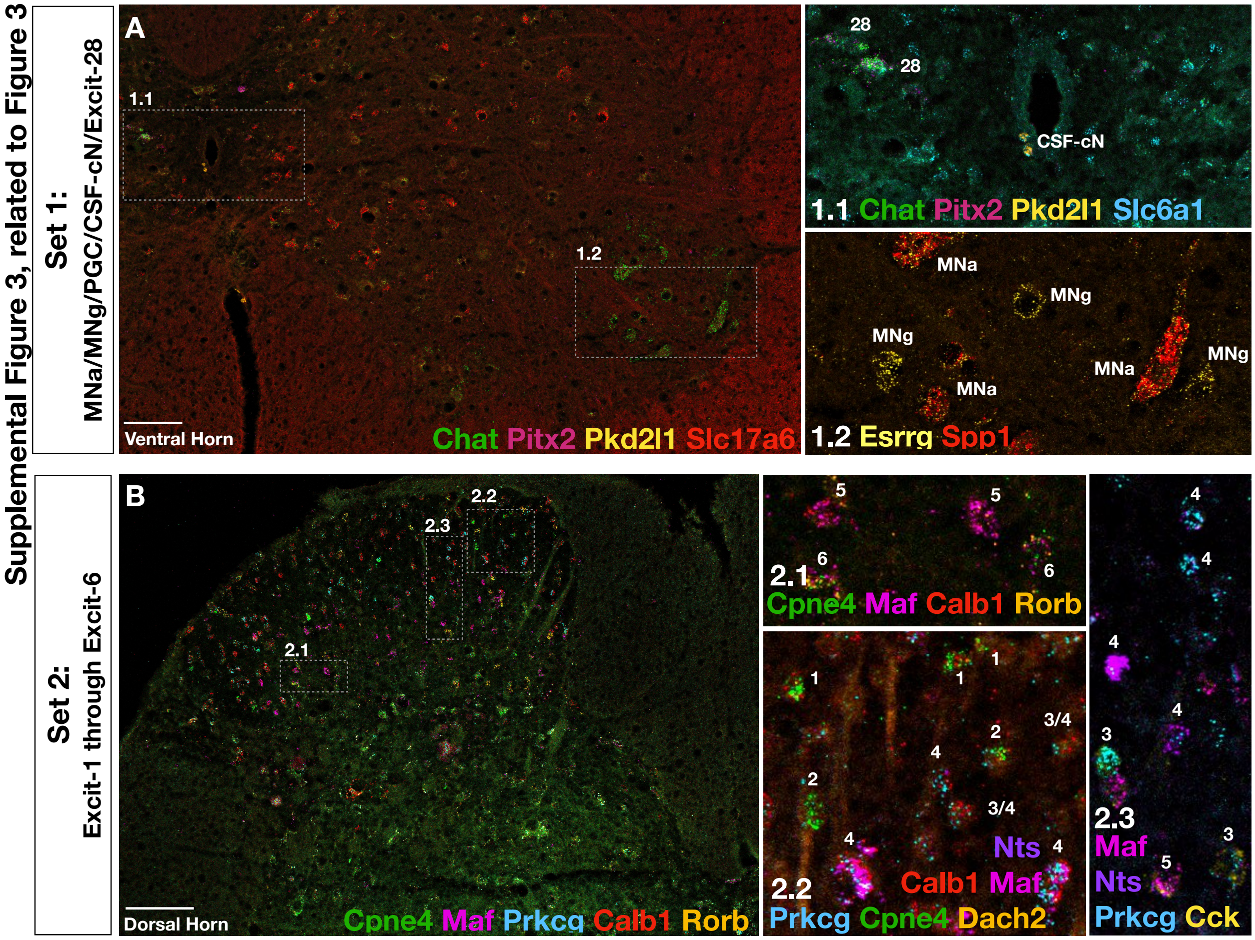


**Supplemental Figure 3 (related to Figure 3). Multi-plexed RNA in situ hybridization of a combinatorial panel of spinal cord cell type marker genes.** For each of the ten sets of RNAScope probes (listed in Supplemental Table 2), this figure shows a 20x tiled image, as well as multiple higher magnification images that are boxed in the 20x tiled image and labeled by Set#.Inset# names. The expressed genes are shown for each image and the cell-type identity is shown by small white numbers next to positive cells in the inset pictures. In many of these images, the contrast and brightness were adjusted using Adobe Photoshop. Scale bars are 100 μm.

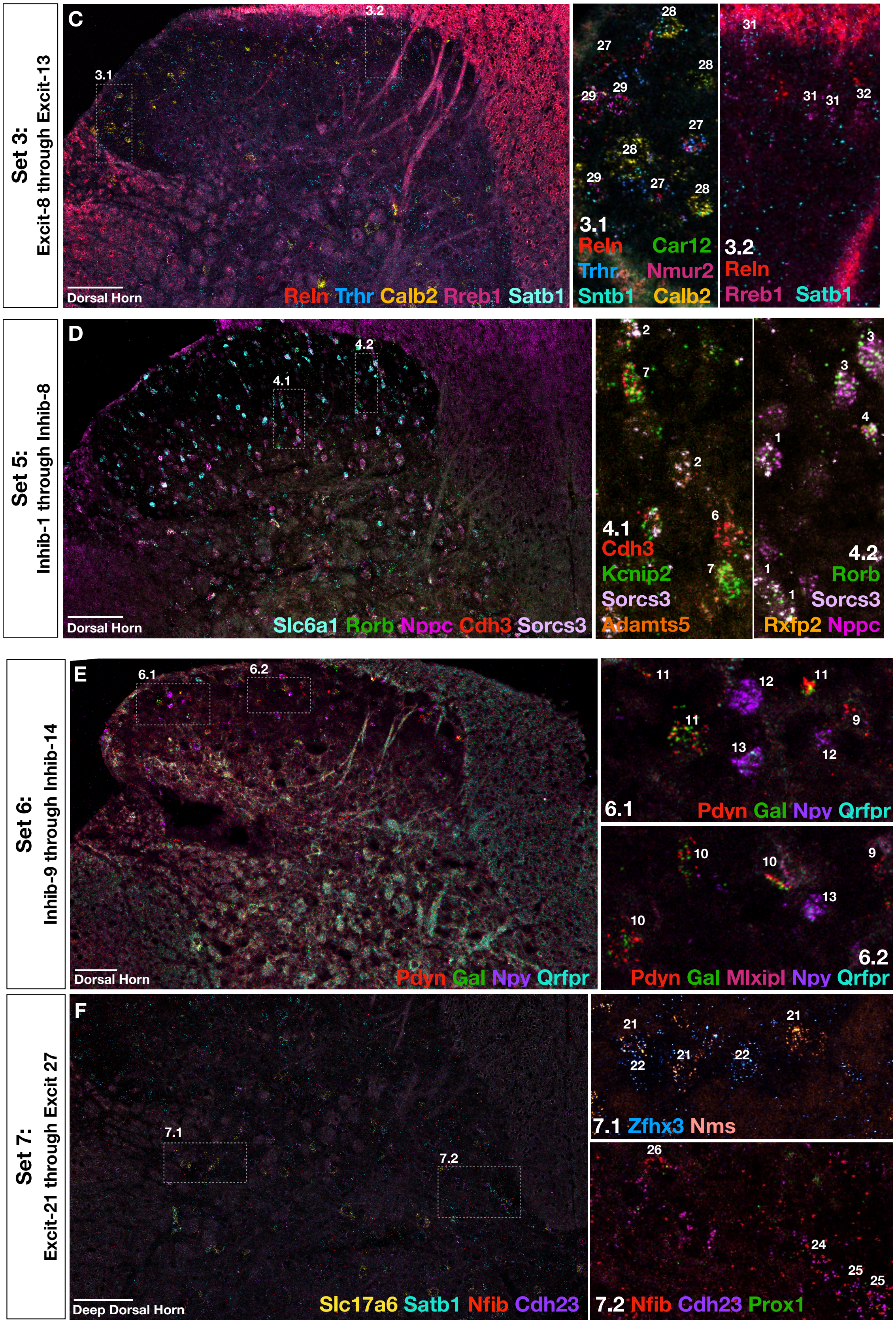
Set 3:
Excit-8 through Excit-13
C
3.1
3.2
Dorsal Horn
Reln Trhr Calb2 Rreb1 Satb1
3.1
Reln Car12
Trhr Nmur2
Sntb1 Calb2
3.2
Reln
Rreb1 Satb1
Set 5:
Inhib-1 through Inhib-8
D
4.1
4.2
Dorsal Horn
Slc6a1 Rorb Nppc Cdh3 Sorcs3
4.1
Cdh3
Kcnip2
Sorcs3
Adamts5
4.2
Rorb
Sorcs3
Rxfp2 Nppc
Set 6:
Inhib-9 through Inhib-14
E
6.1
6.2
Dorsal Horn
Pdyn Gal Npy Qrfpr
6.1
Pdyn Gal Npy Qrfpr
6.2
Pdyn Gal Mlxipl Npy Qrfpr
Set 7:
Excit-21 through Excit 27
F
7.1
7.2
Deep Dorsal Horn
Slc17a6 Satb1 Nfib Cdh23
7.1 Zfhx3 Nms
7.2 Nfib Cdh23 Prox1

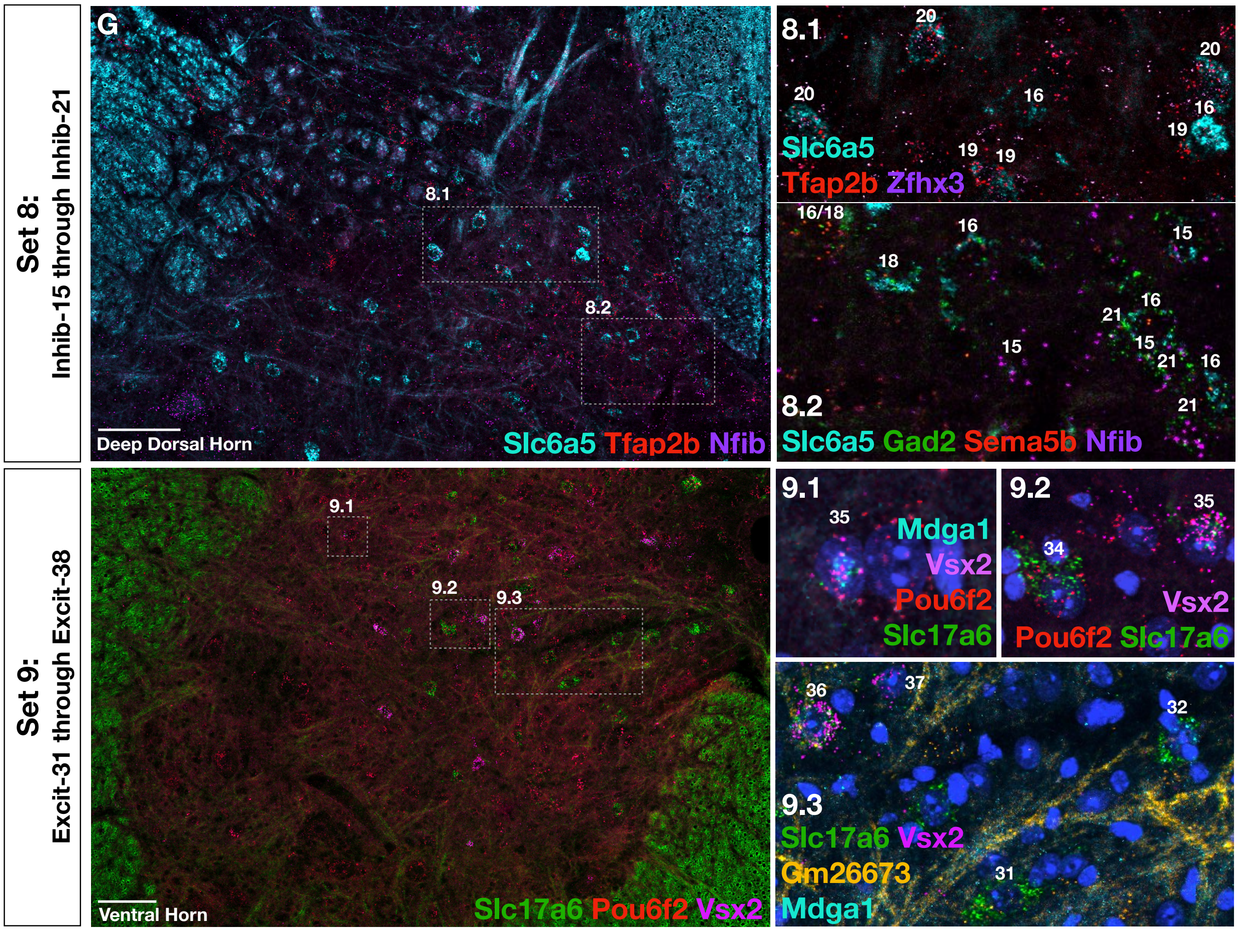

Set 8:
Inhib-15 through Inhib-21
G
8.1
8.2
Deep Dorsal Horn
Slc6a5 Tfap2b Nfib
8.1
20
20
20
16
16
19
19
19
Slc6a5
Tfap2b Zfhx3
16/18
16
15
18
16
21
15
21
16
15
21
8.2
Slc6a5 Gad2 Sema5b Nfib
Set 9:
Excit-31 through Excit-38
9.1
9.2
9.3
Ventral Horn
Slc17a6 Pou6f2 Vsx2
9.1
35
Mdga1
Vsx2
Pou6f2
Slc17a6
9.2
35
34
Vsx2
Pou6f2 Slc17a6
36
37
32
9.3
Slc17a6 Vsx2
Gm26673
31
Mdga1

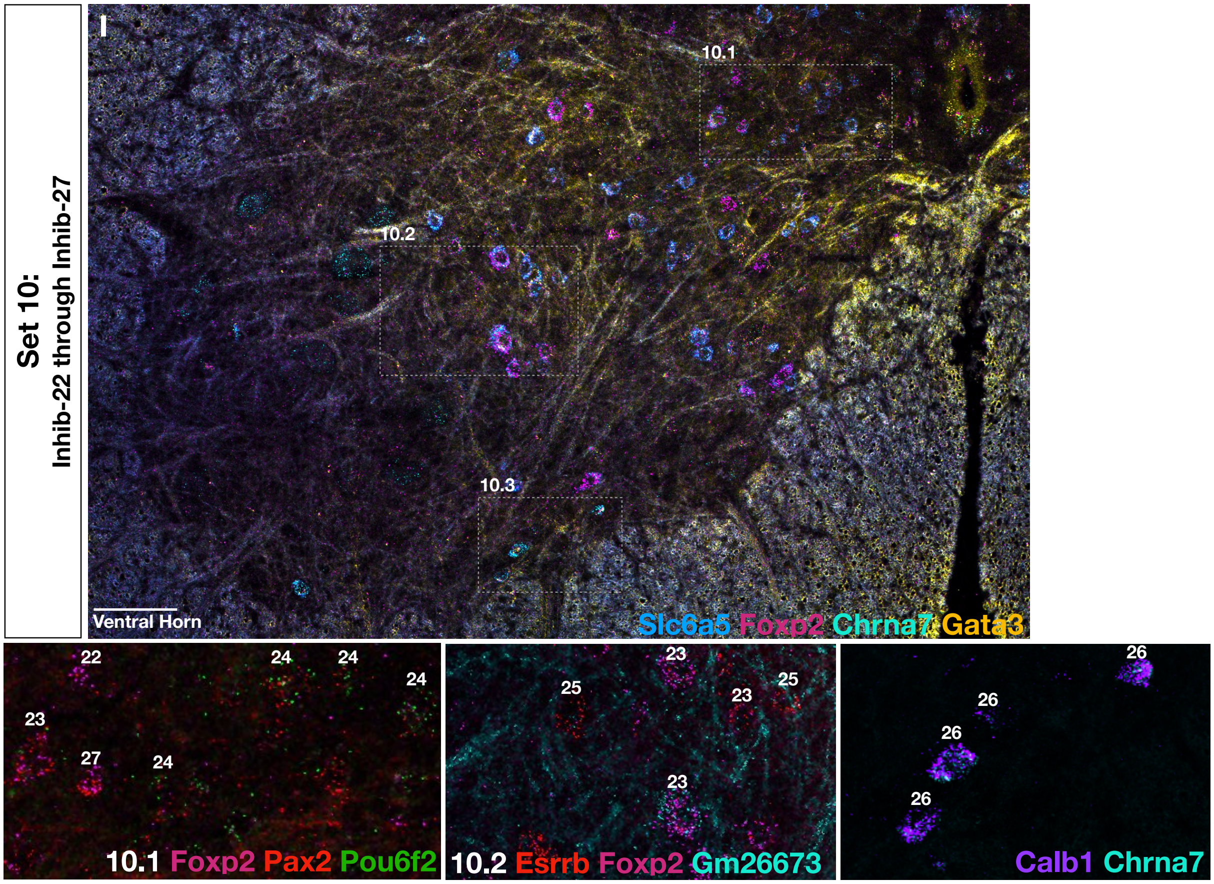
Set 10:
Inhib-22 through Inhib-27
I
10.1
10.2
10.3
Ventral Horn
Slc6a5 Foxp2 Chrna7 Gata3
22
24
24
24
23
27
24
10.1 Foxp2 Pax2 Pou6f2
23
25
23
25
23
10.2 Esrrb Foxp2 Gm26673
26
26
26
26
Calb1 Chrna7

**Supplemental Figure 4 part 1 - related to Figure 5B (plus see next page for 5E Supplemental)**

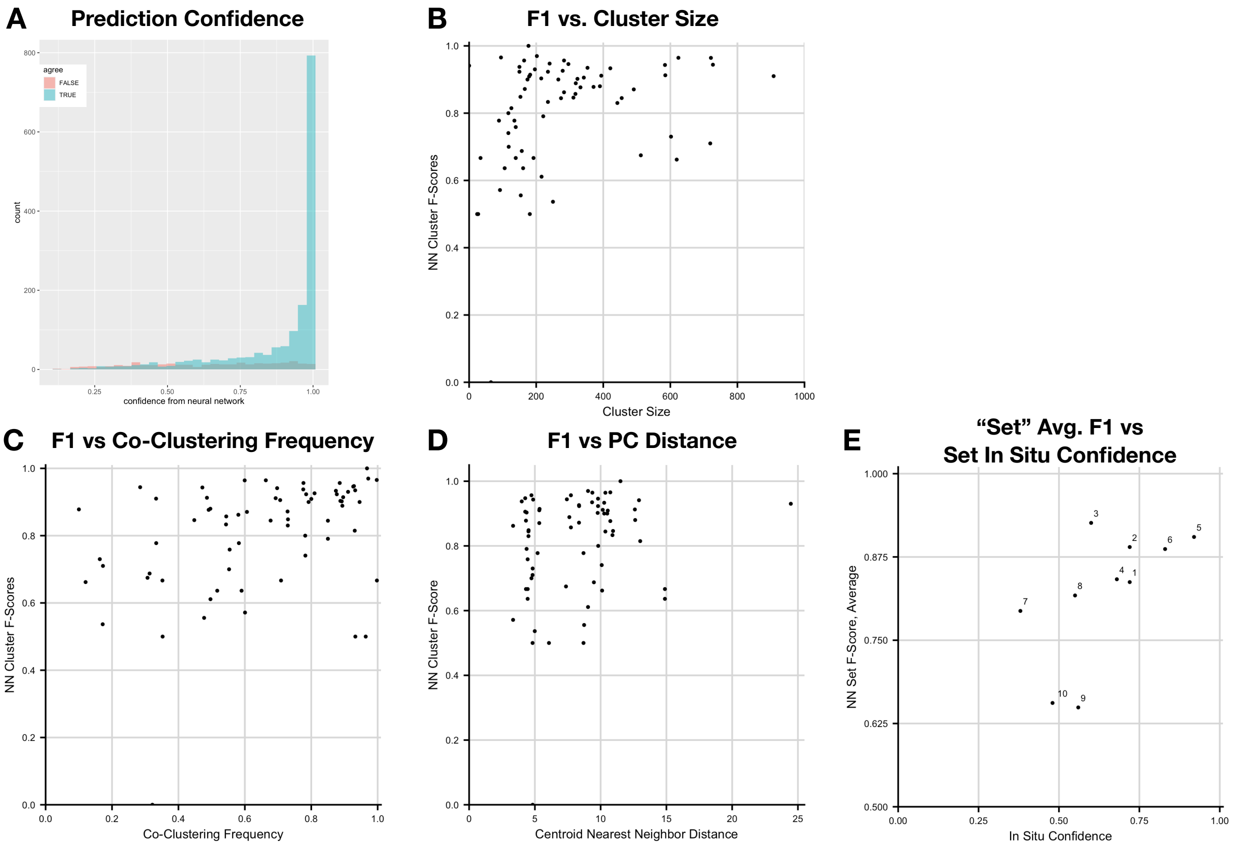


**Supplemental Figure 4. Neural network performance and comparison to cluster robustness; A-E related to Figure 5B; F-I related to Figure 5E.** (A) The confidence (x-axis) with which each cell/nucleus was classified (counts, y-axis), colored by whether the prediction was correct (blue, true) or incorrect (false, pink). (B) Scatterplot of the neural network F1 score and the size of each cluster. (C) Scatterplot of the neural network F1 score and the co-clustering frequency of each cluster. (D) Scatterplot of the neural network F1 score and the distance between the centroid of each cluster and the centroid of its nearest neighbor in 50-dimensional principal component (PC) space. (E) Scatterplot of the neural network F1 score (presented as an average for each "set" of clusters) and the percent of cells in that set that could be confidently assigned to a single cluster by in situ hybridization analysis.

### Supplemental Figure 4 part 2 - related to Figure 5E

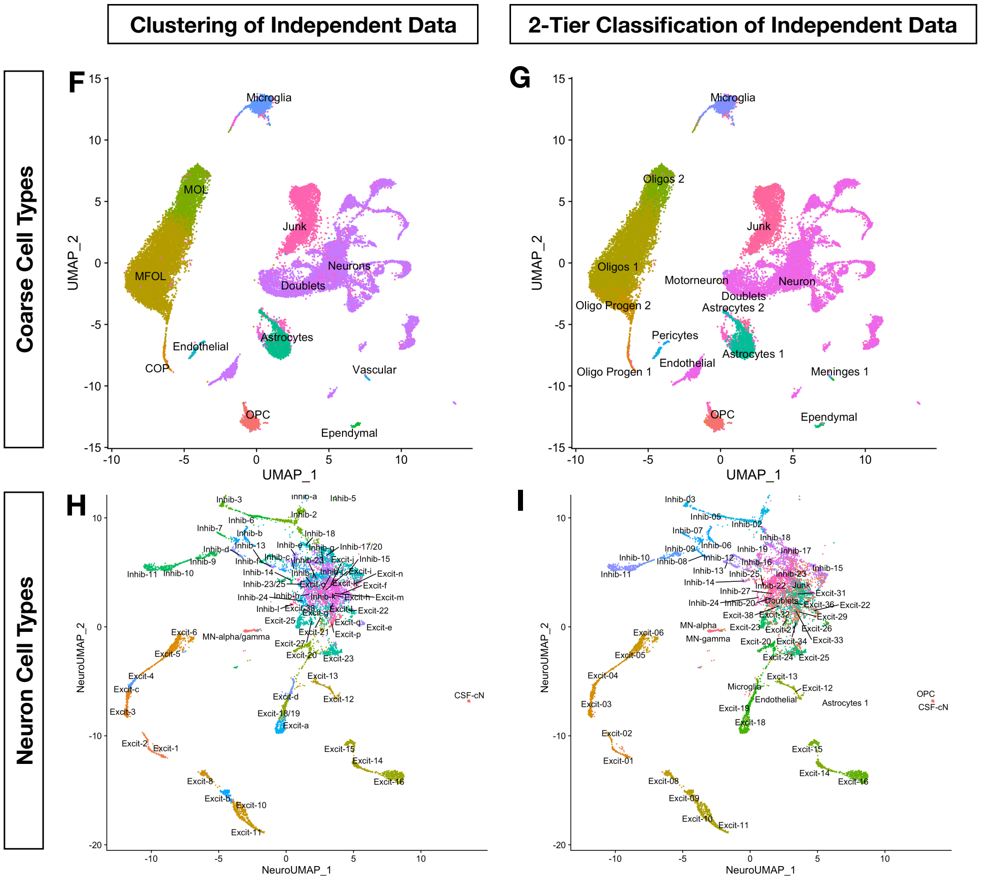


**Supplemental Figure 4. Neural network performance and comparison to cluster robustness; A-E related to Figure 5B; F-I related to Figure 5E.** (F) UMAP plot showing the coarse cell type of each nucleus based on clustering of the independent dataset. (G) UMAP plot showing the coarse cell type of each nucleus based on based on two-tiered computational classification (H). UMAP plot showing the neuron type of each nucleus based on clustering of the independent dataset. (I) UMAP plot showing the neuron cell type of each nucleus based on based on two-tiered computational classification.

**Supplemental Movie 1 (related to Figure 2). Three-dimensional UMAP presentation of 69 populations of spinal cord neurons.** Available as a separate file.

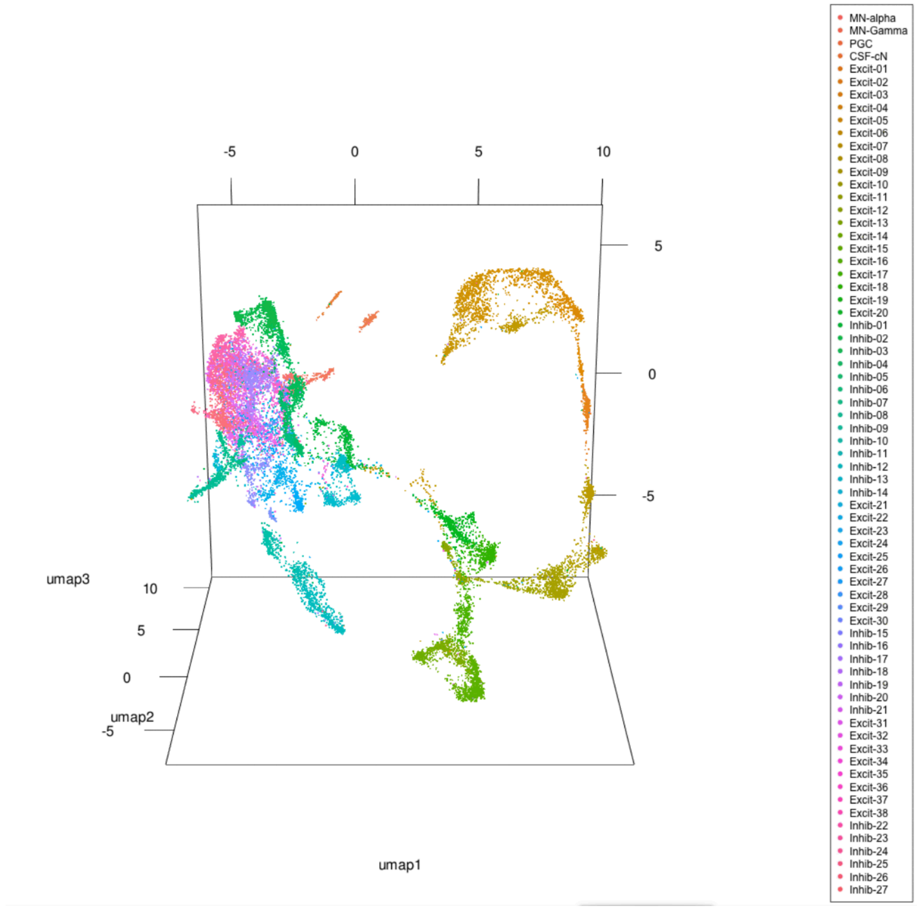

**All tables are available as separate files.**

**Supplemental Table 1 (related to Figures 1 and 2). Top genes associated with each spinal cord cell type (AUROC and Wilcox markers).** The top genes (as sorted by p-value) that are markers for each cluster, compared to all other clusters using AUROC and Wilcox tests.

**Supplemental Table 2 (related to Figure 2). Study contribution for each neuron cell cluster.** The number of neuronal cells/nuclei from each study in each of the 69 neuronal clusters.

**Supplemental Table 3 (related to Figure 3). Combinatorial panel of marker genes tested by in situ hybridization.** (A) The clusters/cell-types that were tested by each set of RNAScope in situ hybridization probes. (B) The genes that were tested in each set. Those that did not work or were not positive in the adult lumbar spinal cord are shown in gray. (C) The number of cells that were counted for each set of probes, with the % of counted cells that could be confidently assigned to one particular cell type from that set or two one of two particular cell types (mean +/- standard error, based on single 14 µm fresh frozen sections, from 3 animals (2 male/1 female or 2 female/1 male). * denotes that the Shox2 probe did not work as part of the Hiplex testing panel but did work using RNAScope v2.

**Supplemental Table 4 (related to Figure 5). Precision, Recall, and F1 scores for the model comparisons in Figure 5, including the different neural network models.**
